# Automatic annotation of the bHLH gene family in plants

**DOI:** 10.1101/2023.05.02.539087

**Authors:** Corinna Thoben, Boas Pucker

**Affiliations:** Plant Biotechnology and Bioinformatics, Institute of Plant Biology & Braunschweig Integrated Centre of Systems Biology (BRICS), TU Braunschweig, Braunschweig, Germany

**Keywords:** bHLH, *Arabidopsis thaliana*, *Croton tiglium*, *Dioscorea dumetorum*, orthology, phylogeny, annotation

## Abstract

**Background:** The bHLH transcription factor family is named after the basic helix-loop-helix (bHLH) domain that is a characteristic element of their members. Understanding the function and characteristics of this family is important for the examination of a wide range of functions. As the availability of genome sequences and transcriptome assemblies has increased significantly, the need for automated solutions that provide reliable functional annotations is emphasised.

**Results:** A phylogenetic approach was adapted for the automatic identification and functional annotation of the bHLH transcription factor family. The bHLH_annotator for the automated functional annotation of bHLHs was implemented in Python3. Sequences of bHLHs described in literature were collected to represent the full diversity of bHLH sequences.

Previously described orthologs form the basis for the functional annotation assignment to candidates which are also screened for bHLH-specific motifs. The pipeline was successfully deployed on the two *Arabidopsis thaliana* accessions Col-0 and Nd-1, the monocot species *Dioscorea dumetorum*, and a transcriptome assembly of *Croton tiglium*.

Depending on the applied search parameters for the initial candidates in the pipeline, species-specific candidates or members of the bHLH family which experienced domain loss can be identified.

**Conclusions:** The bHLH_annotator allows a detailed and systematic investigation of the bHLH family in land plant species and classifies candidates based on bHLH-specific characteristics, which distinguishes the pipeline from other established functional annotation tools. This provides the basis for the functional annotation of the bHLH family in land plants and the systematic examination of a wide range of functions regulated by this transcription factor family.

## Introduction

A basic helix-loop-helix (bHLH) domain is the characteristic and name-giving feature of the bHLH transcription factor family. This gene family is found in three major eukaryotic lineages (animals, plants, fungi). In plants, the bHLH family is one of the largest groups of transcription factors only second to MYBs [1]. To modulate gene expression, the bHLH transcription factors bind as dimers to specific DNA sequences. This function is conserved through two functionally distinct regions in the basic helix-loop-helix domain. The basic region is located at the N-terminus of the domain and consists of mainly basic residues [2]. It functions as a DNA binding domain [3], recognizing a hexanucleotide motif in the major groove [4–6]. At the C-terminus of the domain, the helix-loop-helix region is located. It consists of two amphipathic helices separated by a loop [2]. Through the interaction of hydrophobic residues, it mediates protein-protein interactions used for dimerization [4–6]. Both the formation of homo- and heterodimers was reported for bHLH transcription factors [4, 5, 7]. The most recognized target sequence is the E-box CANNTG [8–12]. While recognizing the E-box, each monomer of the bHLH dimer binds one half of the motif in the major groove [5, 6].

In addition, the bHLHs are also able to interact with other classes of transcription factors [1]. This enables the formation of multimeric complexes [13, 14]. A famous example is the MYB-bHLH-WD40 (MBW) complex, which is involved in the regulation of anthocyanins and proanthocyanidins biosynthesis in the flavonoid pathway and controls epidermal cell fates like trichome initiation or root hair formation [15–18]. The ternary protein complex is composed of a R2R3-MYB, a bHLH, and a WD40-repeat protein [18]. R2R3-MYBs from the subfamilies 5, 6 and 15 and bHLHs from the subfamily 3f can participate in the complex formation. The bHLHs from the subfamily 3f are associated with a conserved N-terminal motif outside the bHLH domain [19].

The functionality of the different elements of a bHLH is determined by certain amino acid residues, which form a conserved motif (Figure 1).

**Figure 1:**
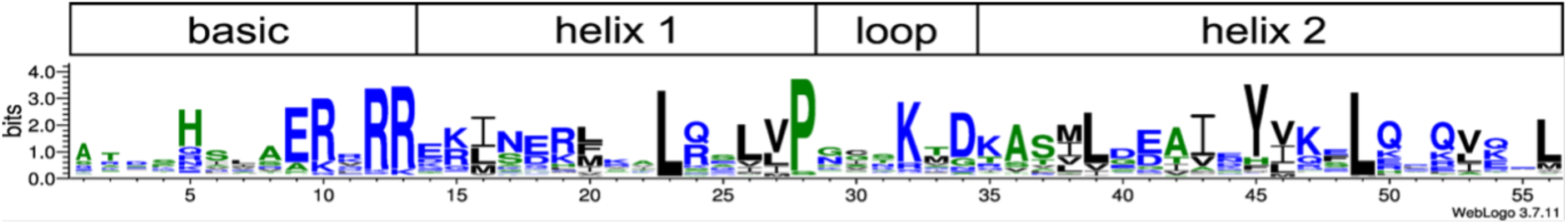
Conservation of amino acid residues of plant bHLH domain. Hydrophobic amino acids are coloured black. Hydrophilic amino acids are coloured blue and neutral amino acids green. The motif is based on conserved residues of bHLHs in Arabidopsis thaliana, Oryza sativa, Physcomitrella patens and Selaginella moellendorffii [10, 11]. Constructed with Weblogo [20].

In plants, the basic region of a bHLH is represented by the first 10 to 19 amino acids [1, 9–12, 21]. Conserved positions in this region determine the DNA binding behaviour [4–6, 22]. Crucial are the positions 5, 9 and 13 in the conserved motif (Figure 1), which mainly define the hexanucleotide target motif [4–6, 22]. The positions 10 and 12 are occupied by arginine residues contacting the DNA backbone [4, 22].

In the helix regions, the hydrophobic amino acids form the core at dimer formation, thus stabilising the interaction [4]. Consistent with this function, analyses of the animal bHLH residues have shown less sequence variability for buried helix positions than for exposed [23]. The loop region shows high variation regarding length and sequence [1, 9–12, 21]. In eukaryotes, the minimum found loop length is five amino acids long [4, 23]. Conserved positions in the loop have been shown to stabilise the shape of the loop [4, 5]. Also, for animal bHLHs, a lysine residue in the loop was reported to participate in DNA binding by interacting with the DNA backbone [4] or being mandatory for DNA binding [24].

Based on their DNA binding properties, members of the bHLH family can be categorised into groups. The first distinction is between DNA binding bHLHs and non-binding HLHs. The latter group is also called ‘atypical bHLHs’. For the prediction of DNA binding ability, the number of basic residues in the basic region is inspected [9]. Most studies use a minimum of 5 basic residues as cutoff to consider candidates as binding bHLHs [11, 12, 25]. The DNA binding bHLHs are further differentiated based on their hexanucleotide target motif. The amino acid composition at specific positions in the basic region determines the motif that is recognized by a bHLH [9]. The most abundant motif in plants and animals is the E-box CANNTG [8–12]. E-box binders are distinguished from non-E-box binders based on the presence of glutamic acid and arginine at the positions 9 and 12 [9, 11, 12, 25–27]. The glutamic acid contacts the first two bases (CA) of the E-box [4–6, 22] and has been shown to be important for DNA binding [28]. This interaction is stabilised by the arginine residue, which directs the side chain of the glutamic acid while contacting the DNA backbone [5]. In plants, as well as animals, the most common E-box motif is the G-box CACGTG [8–12]. Therefore, the group of E-box binders is categorised into G-box binders and non-G-box binders. G-box binders are identified by histidine, glutamic acid and arginine at the positions 5, 9, and 13 [9, 11, 12, 25–27]. The histidine residue binds to the last base (G) of the motif [4, 6, 22]. The arginine residue at position 13 distinguishes the G-box from the E-box motif CAGCTG [4, 6]. In terms of DNA binding specificity, the interaction of bHLH and DNA is not limited to the target motif or basic region. Flanking bases outside the target motif have been shown to discriminate binding for certain animal bHLHs [28] and to be recognized by amino acids in the basic region [5, 6, 22]. Also, residues outside the basic region participate in DNA binding. Contacts with the phosphate groups of the DNA backbone have been observed for residues in the loop and helix 2 region [4, 6], like the lysine residue in the loop mentioned before.

The atypical bHLHs are defined by their lack of basic residues in the basic region and therefore predicted not to be able to bind DNA. Instead, HLHs have shown to contain proline residues in the basic region [9]. Because the HLH region and therefore the ability to dimerize is still intact, they are proposed to be negative regulators of DNA binding by forming heterodimers with DNA binding bHLHs [9, 11]. While this behaviour had been found in both plant and animal bHLHs [13, 29], previous studies found no similarity between the plant and animal HLHs [9, 11].

Based on their phylogenetic relationship, the bHLH transcription factor family can be grouped into subfamilies. Depending on the analysed species and criteria, the number of subfamilies varies. Multispecies analyses have predicted 26 [10] or 32 subfamilies, if atypical bHLHs were included in the analysis [11]. Because the deep notes of the phylogenetic trees in the analyses have low statistical support, no strong conclusion about the relationship between the subfamilies should be drawn. The subfamilies themself are highly supported, allowing examination of subfamily specific characteristics and evolutionary relationship [1, 8–11, 30, 31]. The plant subfamilies are proposed to be monophyletic, as they do not cluster together with other eukaryotic bHLHs [9, 10]. Furthermore, they are conserved among different plant species and most families have been shown to be present in early land plants before the evolution of mosses [10]. Subfamily characteristics are the number and positions of introns [1, 9, 11, 31], the predicted protein length, and the position of the domain [1]. Conserved motifs outside the bHLH domain can be associated with individual subfamilies [1, 9–11, 31]. Some motifs can be observed in different subfamilies, but the relative spatial location is subfamily associated [10]. Most subfamilies belong to the same group with respect to their DNA binding properties. Some distant subfamilies share DNA binding properties which suggests that some features might have developed independently in separate lineages [9, 11]. The functional diversification within the subfamilies is variable. Some subfamily members are involved in similar biological processes or have redundant functions, while other subfamilies contain highly functionally specialised members [1, 11].

In plants, the bHLH family has been expanded through gene duplication [1, 9] and domain shuffling [9–11]. Evidence for this theory are the presence of the same conserved motifs in different subfamilies [10, 11], as well as the sequence diversity outside of the bHLH domain [9]. Domain shuffling also has been suggested for animal bHLHs [30, 32]. Further named arguments regarding animal bHLHs are the loss of the basic domain in some subgroups and the spatial variation of the domain [32], which also apply to plant bHLHs [1, 9, 11].

The bHLH transcription factor family controls a wide range of biological processes, which warrants in-depth investigations of functions and characteristics. The availability of plant sequence data has increased significantly with development of new sequencing technologies like next-generation sequencing (NGS) [33] and more recently long read sequencing technologies [34, 35]. While structural annotation can be achieved automatically with the integration of external hints, the functional annotation of predicted genes remains a challenge. This reinforces the need for automated annotation of sequences.

Various approaches for the automated annotation of sequences are deployed. Established tools assign a functional description via sequence similarity to annotated protein sequences or the detection of hidden Markov Model (HMM) profiles. HMM profiles for this annotation method can be retrieved from Pfam [36, 37] which is a comprehensive protein domain profile database. InterProScan5 is a tool that assigns Pfam domains and other annotation ontology terms to given sequences [38]. To provide standardised vocabularies, Gene Ontology (GO) terms [39, 40], the KEGG Orthology (KO) entries from the KEGG database [41] or the plant specific MapMan BIN ontology [42, 43] can be utilised. The automated functional annotation pipeline Blast2GO assigns GO terms to a set of given sequences based on sequence similarity to previously characterised reference sequences [44]. For KO entries, the KEGG Automatic Annotation Server (KAAS) [45] can be deployed, which is conceptually similar to Blast2GO. Mercator4 [43, 46] is a plant specific functional annotation pipeline, assigning MapMan BIN annotations to novel sequences [47]. The input sequences are classified by scanning for BIN specific HMM motifs. Input sequences that cannot be assigned to a MapMan BIN are annotated by performing a BLASTP search against the Swiss-Prot database [43, 48].

Other functional annotation initiatives rely on the concept that orthologs are likely to have the same function [49], thus specifically identifying orthologs. One approach is the identification via clustering, which is performed by OrthoMCL [50, 51]. Another approach is the investigation of the phylogenetic relationship through identification of homologs, the alignment of all sequences, and the construction of a phylogenetic tree. Based on the inferred relationships in the tree, functional predictions and further analysis can be performed [52]. A similar approach was developed for the identification of gene families based on massive collections of transcriptome assemblies [53]. OrthoFinder implements this approach for the identification of orthologs between species [54]. KIPEs can identify enzymes in a pathway based on orthology to previously characterised sequences and downstream inspection of functionally important amino acid residues [55, 56]. SHOOT can identify the ortholog of a given sequence in a collection of sequence data sets that represent a range of species [57]. The MYB_annotator represents a pipeline for the automated functional annotation of the MYB transcription factor family by phylogenetic identification of orthologs and a detailed characterisation of identified candidates based on specific sequence features [58].

Here, the objective was to develop a pipeline for the automated identification and functional annotation of bHLHs in plants based on orthology to previously characterised sequences. Initial candidates are recognized through similarity to a collection of bait sequences or by a bHLH-specific HMM motif. The pipeline harnesses a phylogenetic approach to define orthologs as final candidates. Special characteristics of the bHLH family like the subfamily-specific motifs and DNA binding properties are analysed. Screens of data sets with numerous isoforms like *de novo* transcriptome assemblies are enabled by a parallelization option.

## Materials and methods

### Development of a workflow for automatic bHLH annotation

An automatic basic helix-loop-helix (bHLH) annotation pipeline was implemented in Python3 [55]. The bHLH_annotator builds on a strategy derived from the MYB_annotator [58] and includes the identification of bHLH candidates based on sequence similarity revealed through BLASTp v2.12.0+ [60], the construction of a global alignment of candidate sequences with MAFFT v7 [61] or Muscle5 [62], the construction of phylogenetic trees with FastTree2 v2.1.10 [63] or RAxML-NG v0.9.0 [64], the identification of orthologs and paralog clusters with DendroPy 4.5.2 [65], the domain check and screening for subfamily specific motifs with HMMER 3.3.2 [66] and the prediction of DNA binding group based on residues in the basic region (Figure 4). The MYB_annotator [58] was optimised to annotate members of the MYB family with a highly conserved repeat motif. The bHLH_annotator was adjusted to the highly variable bHLH domain in members of the bHLH family. In contrast to the MYB_annotator [58], Muscle5 [62] is set as the default alignment tool because it provided a more reliable alignment of the bHLHs in our tests. To allow alignments with Muscle5 [62] for a high number of identified candidates (over 600 candidates), the bHLH_annotator was equipped with an optimised bait collection and a parallelization option, which reduces the computational costs of the classification step by partitioning the candidates into bins that are separately analysed. Furthermore, additional steps analysing bHLH specific characteristics were added. They include a domain check and extraction of the bHLH domain, screening for subfamily specific motifs with HMMER 3.3.2 [66], and the prediction of DNA binding groups based on residues in the basic region (Figure 4).

### Input data sets

If run with default settings, the reference sequences used to annotate ortholog candidates are the *Arabidopsis thaliana* bHLHs. These sequences were annotated by the latest public data release on The Arabidopsis Information Resource (TAIR) website [67, 68]. Also, the subfamily name of each reference sequence was retrieved from a multi species study of the bHLH family [10]. From the same study [10], sequences of subfamily specific motifs were collected, aligned with Muscle5 [62], and a HMM motif specific for a subfamily was built with the HMMER 3.3.2 [66] function “hmmbuild” if possible. For the prediction of DNA binding groups, AT1G09530 was chosen as reference. Residue positions of the candidate bHLH domains are determined based on this polypeptide sequence.

### Bait sequence collection

Previously described bHLH sequences from various plant species were retrieved to form the bait sequence collection. Sequences were taken from the supplementary data of the respective publication or extracted from the plants predicted polypeptide sequences specified in the respective publication (Table 1). Duplicate sequences were removed from the bait collection. bHLHs with a high grade of sequence divergence were identified in the well studied species *A. thaliana* [1, 9, 11] and *Oryza sativa* [11, 31] (see Additional file 1 for details). The BLASTp-based Python script “collect_best_BLAST_hits.py” in version v0.29 [69] was used to find the best BLAST hits in predicted polypeptide sequences of *Brassica napus* for the diverged sequences in *A. thaliana* and *Zea mays* in *O. sativa.* The obtained sequences were added to the bait sequence collection to represent the diverged sequences with a higher phylogenetic diversity.

**Table 1:**
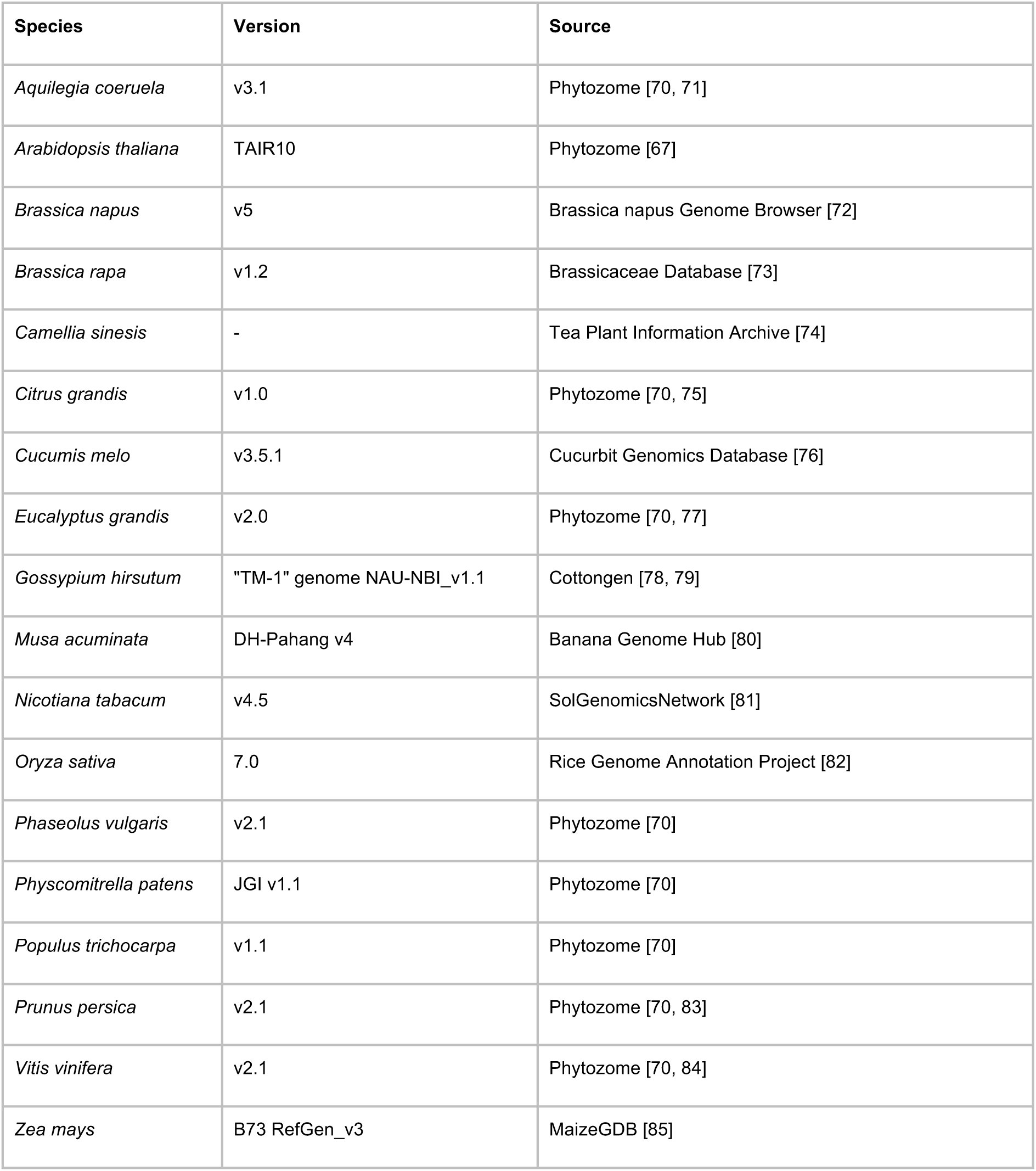
Plant polypeptide sequences used to collect bHLH sequences. The species, version and database source are given.

Additional bHLH-like outgroup sequences were identified based on sequence similarity to bHLH sequences and a phylogenetic placement outside of the monophyletic bHLH clade (see Additional file 2 for details). The collection of bait and outgroup sequences was reduced to represent the full sequence diversity with a minimal number of sequences. This optimisation of the bait sequence collection substantially reduces the computational costs and run time of the analysis (see Additional file 2 for details). With the HMMER 3.3.2 [66] option “hmmbuild” an HMM motif of the optimised bait collection was created.

The bHLH bait sequence collection and outgroup sequence collection represent the bait collection v0, which was used for benchmarking. For the bait collection v1, the bHLH candidates (type 1 and 2) identified in the benchmarking of the *A. thaliana* Col-0 accession and not described in the literature [1, 9, 11] were added. Also, the remaining bHLH sequences of *B. napus* and *Z. mays* were added, as these species are only represented by the diverged sequences in v0. The final bait collection v1.1 was obtained by removing bait sequences with an interchanged phylogenetic placement regarding the bHLH baits and outgroup.

A phylogenetic analysis of the bait collection v1.1 and the optimised bait collection v1.1 was performed to identify subfamilies proposed by previous multi-species studies [10]. For each subfamily, a Weblogo was created and the representation of the land plant lineages in the subfamilies was analysed (see Additional file 3 for details).

### Parameter optimisation

Optimal BLAST parameters would allow a comprehensive identification of all bHLHs while minimising the number of false-positive candidates that need to be filtered out in the following steps. To avoid overfitting of parameters, the optimisation was performed on *A. thaliana* as representative of eudicot plants as well as *O. sativa* as representative of monocot plants (see Additional file 1 for details).

### Proof of concept and benchmarking

As proof of concept, the pipeline in v1.01 with the bait collection v0 was deployed on the Araport11 polypeptide sequence collection of the *A. thaliana* accession Col-0 [68]. *A. thaliana* bait sequences were temporarily removed from the bait sequence set before deployment. Afterwards, the pipeline was deployed on polypeptide sequences of the *A. thaliana* accession Nd-1 [86], *Dioscorea dumetorum* [87] and *Croton tiglium* [88] with the complete bait collection. The identified candidates in the *A. thaliana* Nd-1 accession were assigned to the Col-0 candidates by using them as baits for the BLASTp-based Python script “collect_best_BLAST_hits.py” v0.29 [69] to compare the results of both accessions.

## Results

### bHLH and outgroup bait collection

The initial bait collection consists of 4,545 bHLH sequences of 28 plant species [89] (Additional file 4, Additional file 5). The optimisation process resulted in an optimised bait collection of 318 sequences representing 27 species (Additional file 6). For each sequence the species name, the reference labelling it as bHLH, the accession identifier, and the source of the sequence was documented. As representatives of bryophytes, lycophytes, gymnosperm, and several angiosperm species are included, both the bHLH bait collection and the optimised bait collection cover a wide phylogenetic range of land plant species (Figure 2). The collection of outgroup sequences contains 136 identified non-bHLH sequences of 16 species (Additional file 7). The optimised outgroup collection contains 84 non-bHLHs and still represents the same 16 species (Additional file 8).

**Figure 2:**
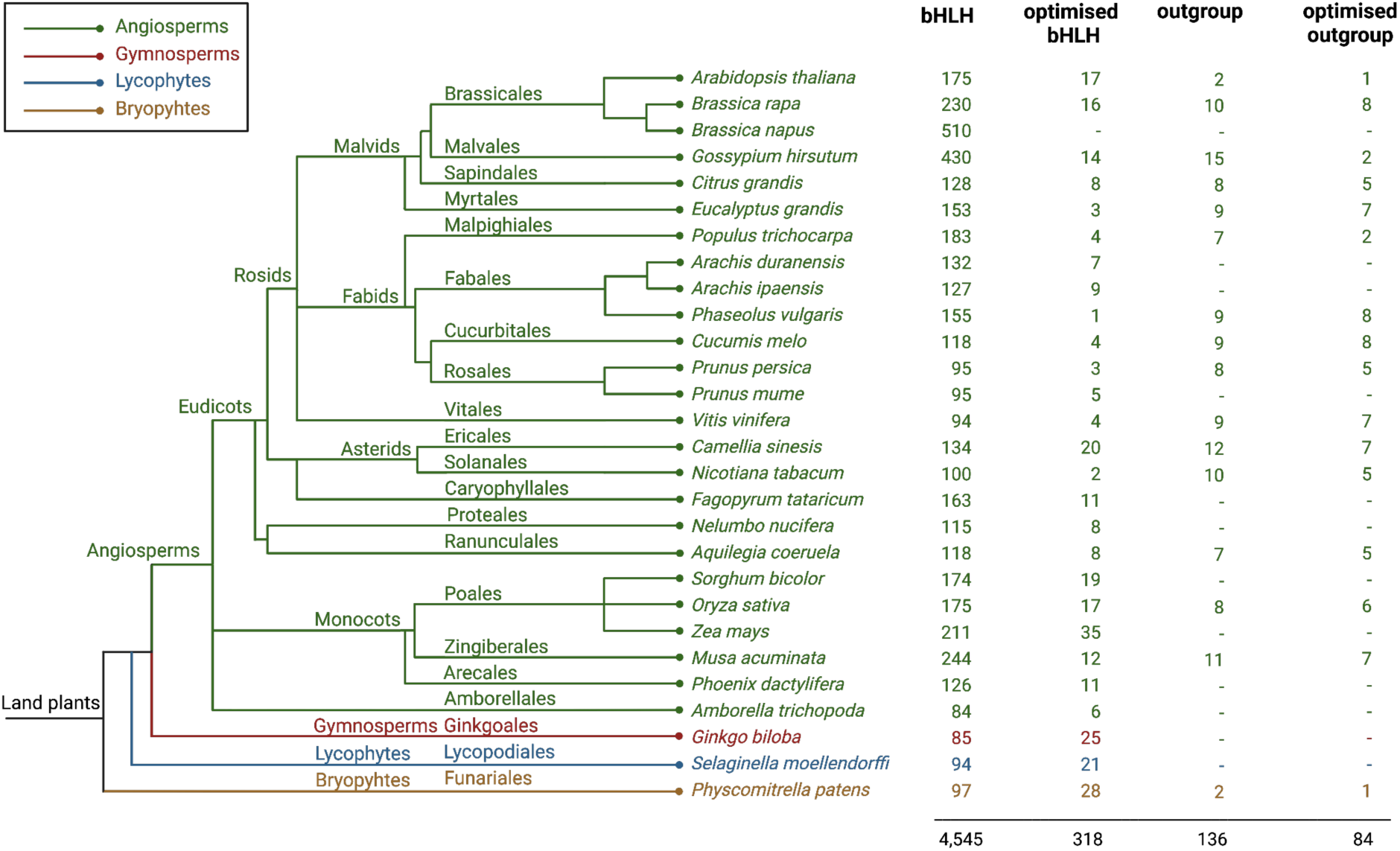
Phylogenetic relationship of all plant species represented in the bait sequence collection v1.1. Number of sequences in bait collection, optimised bait collection, outgroup collection and optimised outgroup collection. Different plant lineages are indicated with colours: Angiosperms (green), Gymnosperms (red), Lycophytes (blue), and Bryophytes (yellow). Orders of plant species were retrieved from WFO [90]. Phylogenetic relationship of orders and families revised by [91, 92].

The phylogenetic analysis identified 27 subfamilies in the bait collection and optimised bait collection (Additional file 3, Additional file 9). The Weblogos of the subfamilies (Additional file 10) demonstrate a variable position of the bHLH domain between the subfamilies, which conforms with the results of previous studies [1, 9–11]. As observed in other multi-species studies [10, 11], the subfamilies are highly conserved among the land plant lineages (Additional file 3, Additional file 11).

The phylogenetic relationship shows the outgroup sequences forming a group separated from the monophyletic bHLH sequences (Figure 3, Additional file 12). The phylogenetic separation of the outgroup sequences is also shown in a phylogenetic tree inferred with IQ-TREE 2 [93] (Additional file 13). A few bHLH sequences of the optimised bait collection are not matched by the HMM motif of the optimised bait collection.

**Figure 3:**
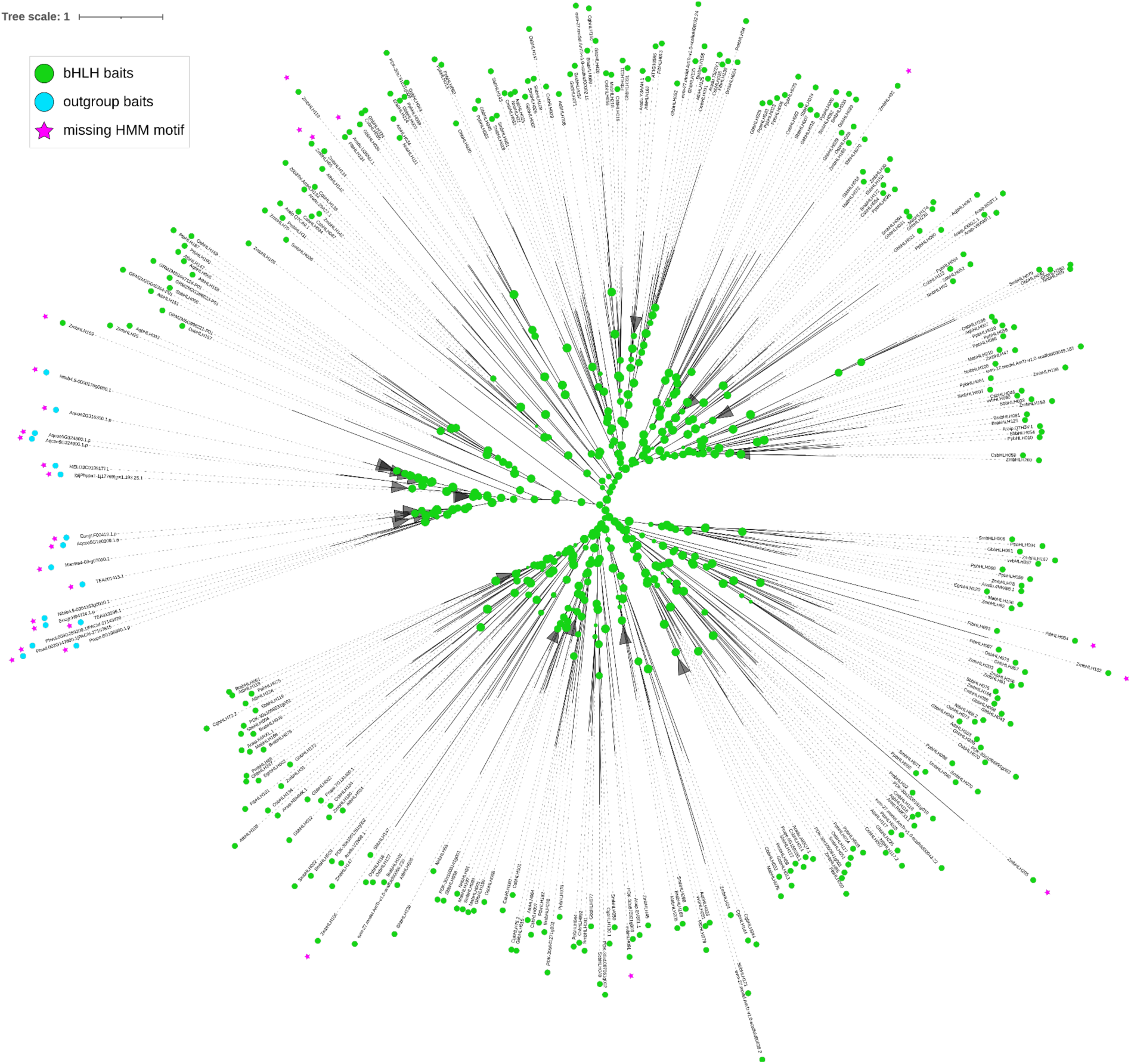
Maximum likelihood tree showing the phylogenetic relationship between optimised bHLH bait collection and optimised outgroup collection v1.1. Bootstrap values are represented by the size of green circles. Clades with an average branch length distance below 0.8 are collapsed. Figure created with iTOL [94].

### bHLH identification and annotation workflow

The pipeline comprises several mandatory and optional steps (Figure 4). In step 1, initial candidates are identified via BLAST [95, 96] or HMMER [66]. In step 2, the candidates are classified as ingroup (bHLH) or outgroup (non-bHLH) members. For a high number of identified candidates (over 600 candidates), a parallelization option is provided in this step. A phylogenetic tree containing the clean candidate and bait sequences is constructed (step 3). Based on the tree, orthologous baits and orthologous bHLH reference sequences are assigned to each candidate (step 4). In the following steps, an analysis of the bHLH domain is performed based on a HMM motif and the DNA binding group is predicted (step 5). Also based on HMM motifs, subfamily specific motifs are identified in the candidate sequences (step 6). Next, phylogenetic trees are constructed containing clean candidate sequences and *A. thaliana* bHLHs (step 7), respectively. Step 8 provides the option to collapse large groups of similar sequences by defining one representative per clade. This option is intended for the analysis of *de novo* transcriptome assemblies, which are usually rich in transcript isoforms. In step 9, another phylogenetic tree of the representative candidates retained in step 8 and *A. thaliana* bHLHs is created. The bHLH_annotator pipeline is publicly available through a GitHub repository: https://github.com/bpucker/bHLH_annotator.

**Figure 4:**
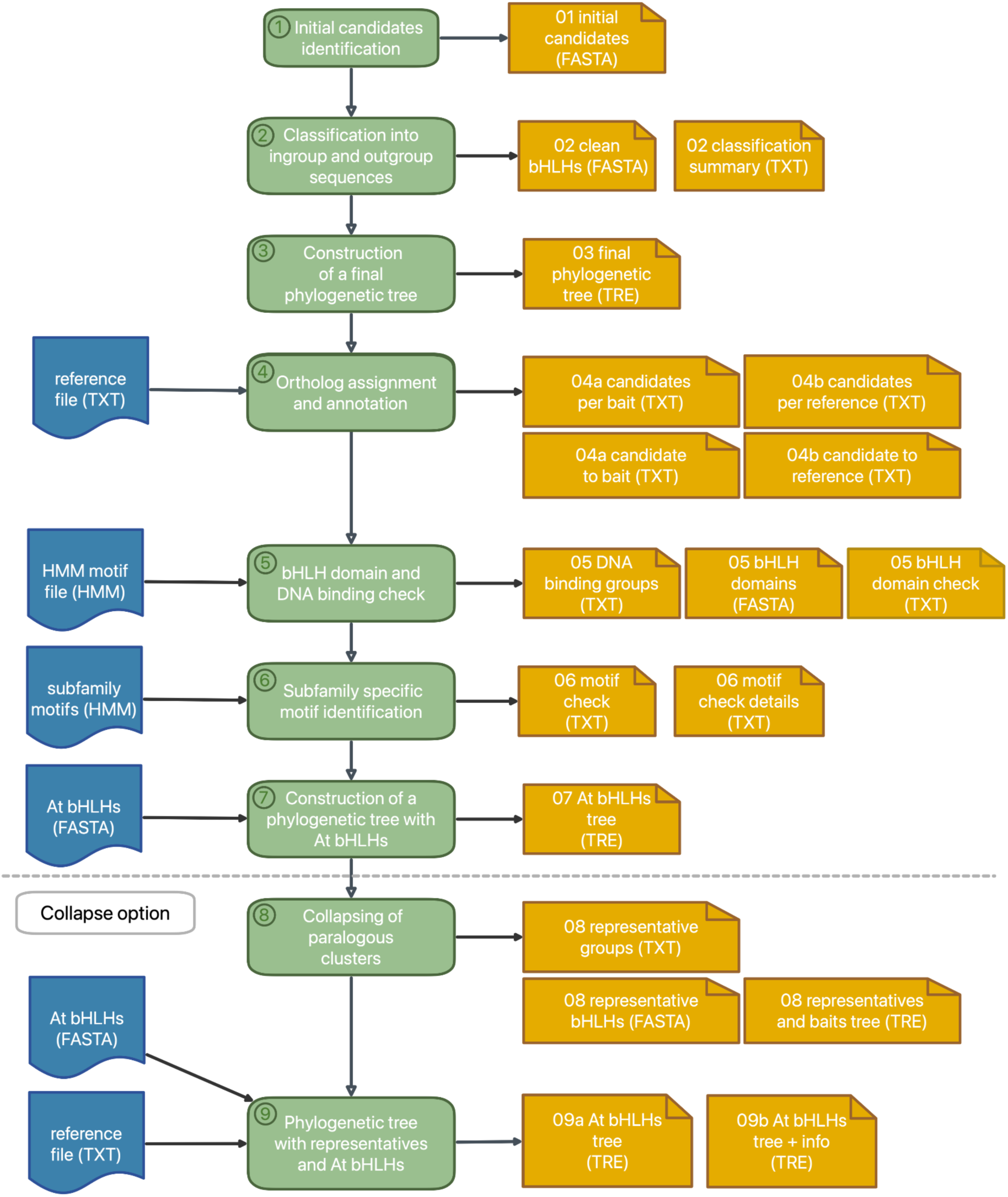
Schematic illustration of the bHLH_annotator pipeline. The pipeline steps and output files are numbered. Input files are coloured blue. The pipeline steps are coloured green. All output files with format are numbered and coloured yellow. Step 8 and step 9 are only performed if the collapse option is chosen.

#### Step 0: Input data and integrity check

Prior to execution of the bHLH_annotator, several inputs are checked. Users need to supply an input FASTA file containing coding or peptide sequences (subject file) and to specify an output folder. Additional input files like the HMM motif for the HMMER search can be defined in a config file or as arguments.

The defined files are checked for forbidden characters in the sequence identifiers and for consistency of identifiers across files. The input FASTA file is cleaned to remove forbidden characters. A mapping file is created to document the connection between cleaned identifiers and the original identifiers of the user-supplied subject file.

#### Step 1: Identification of initial candidates

Initial candidates can be identified via BLAST [95, 96] or HMMER [66]. For HMMER, a HMM motif must be defined (the HMM motif of the bait collection is defined as default in the config file). As default option, BLAST is used to perform a search with the bait sequence collection against the user-supplied subject file. The hits are filtered based on bit score, alignment length and percentage of identical matches (described as similarity) to retain the initial bHLH candidates.

#### Step 2: Classification into ingroup and outgroup sequences

A phylogenetic tree of the initial candidates and bait collection is constructed. As an alignment tool, Muscle5 [62] or MAFFT v7 [61] can be selected with Muscle5 [62] being the default option. Each alignment is trimmed by removing positions with less than 10% occupancy in a given alignment position. Tree construction can be done via default FastTree2 [63] with option “-wag” or RAxML-NG [64]. In case of a high number of candidates (over 600 candidates), parallel tree construction with a fixed number of candidates per tree is possible with the parallelization option. This is recommended as it allows the reduction of computational costs and substantially reduces the run time. All candidates are screened for the presence of the defined HMM motif. The classification of candidates is based on the phylogenetic distance to neighbouring ingroup and outgroup leaves representing bait sequences (Figure 5).

**Figure 5:**
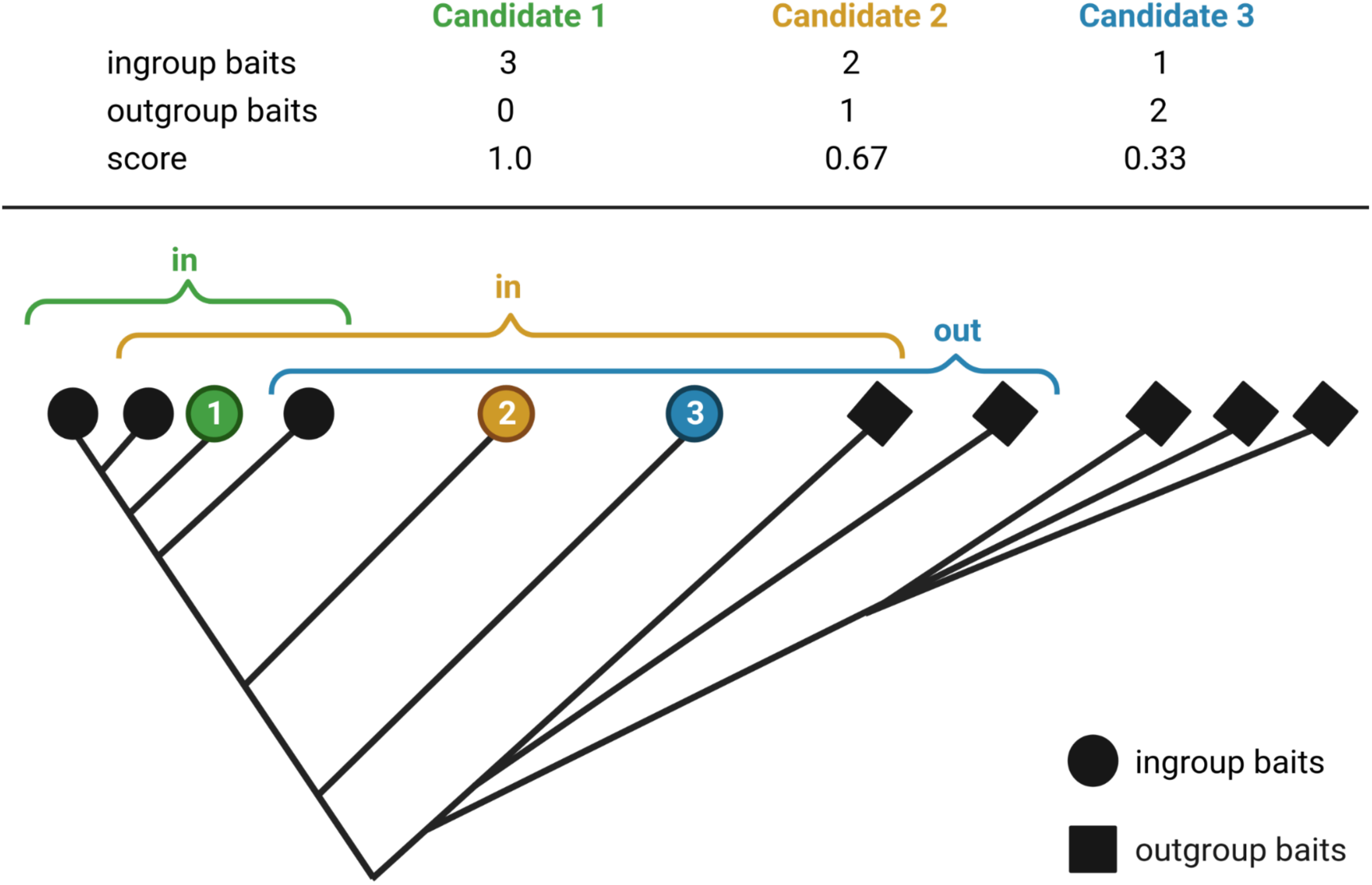
Schematic illustration of the classification into ingroup and outgroup candidates. The brackets denote the neighbouring leaves considered for the classification of each candidate. The number of considered neighbouring leaves for classification is 3. Candidates with a score higher than 0.5 are considered as ingroup bHLHs (in); candidates with a score below 0.5 are considered as outgroup candidates (out).

With DendroPy 4.5.2 [65], patristic distance and edge count of each candidate leaf to all other leaves of the trees are computed. The classification is performed on each candidate leave separately. Neighbouring leaves are sorted based on ascending edge count and a defined number of these leaves is selected for classification. Singular leaves with a patristic distance longer than the mean nearest taxon distance (calculated with DendroPy v4.5.2 [65] multiplied by a defined factor are removed from the selection to exclude outliers. The ingroup and outgroup baits among the selected leaves are counted and the relative contribution of ingroup sequences is calculated as a score. Based on a defined minimum score and a minimum number of ingroup and outgroup baits in the selection, the candidate leaf is classified as ingroup or outgroup. Also, candidates missing the bHLH motif can be excluded from the ingroup if the user specified the presence of this motif as mandatory.

With the candidates classified as ingroup, a second classification with a newly constructed tree is started to filter candidates that were previously falsely accepted. The second phylogenetic tree is expected to be of higher quality due to the reduced number of non-bHLHs in the multiple sequence alignment. The ingroup candidates of the second classification are accepted as final bHLH candidates.

#### Step 3: Construction of a final tree

A phylogenetic tree of the final candidate and bait sequences is constructed as described above. Because the contribution of non-bHLH sequences is the smallest among all analyses, the resulting tree is assumed to represent the phylogenetic relationship of the candidates with the highest possible reliability.

#### Step 4: Assignment of orthologs and annotation

An ortholog bait sequence is assigned to each final bHLH candidate. DendroPy v4.5.2 [65] is used to calculate the edge and patristic distances from each candidate to all baits in the final tree (step 3). The bait sequence with the minimum edge distance to each candidate is defined as ortholog. As a result, all ortholog candidates per bait sequence are collected. Also, for all candidates, the assigned bait sequences are listed together with edge distance and patristic distance.

In the following step, the final bHLH candidates are annotated based on orthologous relationships to the reference sequences. The reference file defines the functional annotation description, an alternative sequence name, and the name of the phylogenetic subfamily for all reference sequences. Based on the assigned orthologs, the candidates are annotated with this information from the reference file. While annotated *A. thaliana* bHLHs are provided as standard references, users could also run the analysis with their own references. The reference sequences must be included in the complete bait collection, or the *A. thaliana* bHLHs file (step 7).

#### Step 5: bHLH domain check and prediction of binding group

All final candidates are checked for the presence of the bHLH domain using HMMER [66] and the defined HMM motif (the HMM motif of the bait collection is defined as default in the config file). The results are summarised and identified bHLH domains are exported into a FASTA file. For prediction of DNA binding groups, the crucial domain positions are analysed for each candidate. Based on determined positions in the reference bHLH AT1G09530, the crucial locations are identified in the trimmed alignment that served as basis for the final phylogenetic tree (step 3). For each candidate, several information are listed: (1) the number of basic residues in the basic region, (2) the amino acids at the positions of 5, 9 and 13 (based on AT1G09530), (3) DNA binding ability, (4) E-box binding ability, and (5) G-box binding ability.

#### Step 6: Identification of motifs

All final candidates are screened for the presence of subfamily specific motifs using HMMER [66]. For each candidate, the recognized motifs are listed, and the motif sequences are extracted. The subfamily specific motifs are defined in the motifs file and the collection can be extended by users.

#### Step 7: Construction of a tree with *A. thaliana* bHLHs

A phylogenetic tree of the final candidates and *A. thaliana* bHLHs is constructed as described above. While most users might consider the well-studied *A. thaliana* bHLHs helpful, it is also possible to provide the bHLHs sequences of a different species or bHLH sequences of multiple species through this option.

#### Step 8: Collapsing paralogous clusters

*De novo* transcriptome assemblies are characterised by a high level of sequence redundancy due to large numbers of transcript isoforms per gene. Large groups of similar sequences (expected paralogs or isoforms) are collapsed by defining one representative sequence per clade. For each leaf of the final tree (step 3), the mean nearest taxon distance and patristic distance to all other leaves of the trees is calculated using DendroPy v4.5.2 [65]. The neighbouring leaves are sorted by ascending edge distance and are analysed in this order. All candidate leaves with a patristic distance less than the mean nearest taxon distance multiplied by a defined cutoff factor are identified as members of the paralog group. The analysis is stopped at the first bait sequence. All members of the paralog group are excluded from further group member identifications to prevent overlapping clusters. Each paralog is collapsed by defining the member with the longest sequence as representative. A phylogenetic tree of the representatives and bait sequences is constructed under the same conditions as described above.

#### Step 9: Construction of phylogenetic tree with representatives and *A. thaliana* bHLHs

A phylogenetic tree of the representative candidates and the *A. thaliana* or other landmark bHLH sequences is constructed as described above (step 7). Also, another phylogenetic tree is constructed containing the subfamily and alternative sequence names of all reference bHLH sequences in their labels. Based on the phylogenetic relationship to the references, subfamilies are assigned to the representative candidate sequences.

### Proof of concept and benchmarking

#### *A. thaliana* accession Col-0

As a technical validation, the Col-0 sequences were screened for bHLHs. The pipeline identified 571 initial candidates in the *A. thaliana* accession Col-0 [68]. After the first phylogenetic classification in the pipeline, this number was reduced to 249 candidates. The second classification identified 241 possible bHLH candidates (Figure 6), as eight candidates were placed differently in the phylogenetic tree of the more stringent second classification round.

**Figure 6:**
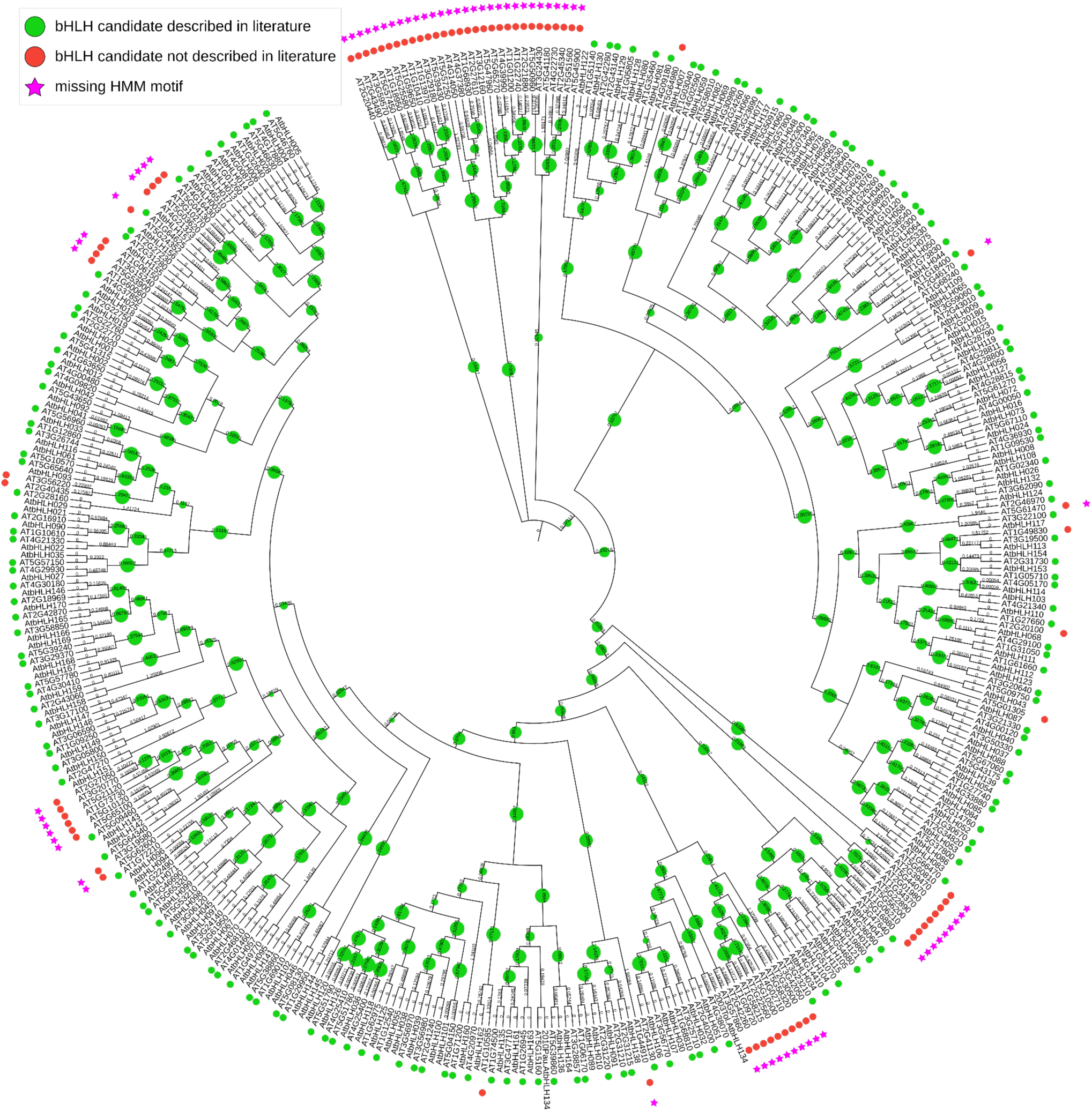
Cladogram derived from a maximum likelihood tree showing the phylogenetic relationship between identified candidates in A. thaliana accession Col-0 and A. thaliana bHLHs of the bait collection. Bootstrap values are represented by green circle size. Figure created with iTOL [94].

These possible bHLH candidates included 165 of the 167 *A. thaliana* bHLHs that were described in the literature [1, 9, 11]. The two candidates not identified are AT1G25310 and AT5G50010, which were not detected by BLAST due to a sequence similarity below 40% and/or bit score below 60. The 76 candidates identified in addition to previous reports in the literature can be divided into four types: **(1)** 5 bHLH candidates that are not mentioned in publications [1, 9, 11], but are annotated as bHLHs in TAIR; **(2)** 3 sequences not annotated as bHLH but harbouring the bHLH domain and placed within bHLH clades; **(3)** 13 candidates missing the bHLH motif and not being annotated as bHLH, but placed within bHLH clades; (**4)** 55 outliers on singular long branches or in groups on long branches with not more than one bait sequence. With the filter option to exclude candidates missing the bHLH motif, the false positives were reduced to members of the first type (AT1G10585, AT1G06150, AT5G01305, AT2G20100, AT1G49830) i.e. bHLHs not mentioned in the publication, but annotated as bHLH in TAIR and the second type (AT2G40435, AT3G56220 and AT5G64980) i.e. not annotated as bHLH but harbouring the characteristic domain.

When deploying the pipeline with the option to search for initial candidates via HMMER [66] instead of BLAST [95, 96], 198 initial candidates were identified. The classification reduced this number to 186 possible bHLH candidates. These candidates included all 167 *A. thaliana* bHLHs of the bait collection and 19 additional candidates. The additional candidates included all members of type one and two identified in the BLAST search.

#### *A. thaliana* accession Nd-1

To test the pipeline on a well-defined annotation without perfectly matching sequences, another *A. thaliana* accession was analysed. The pipeline identified 584 initial candidates in the *A. thaliana* accession Nd-1 [86]. After the first classification, this number was reduced to 239 candidates. The second classification identified 235 bHLH candidates (Figure 7).

**Figure 7:**
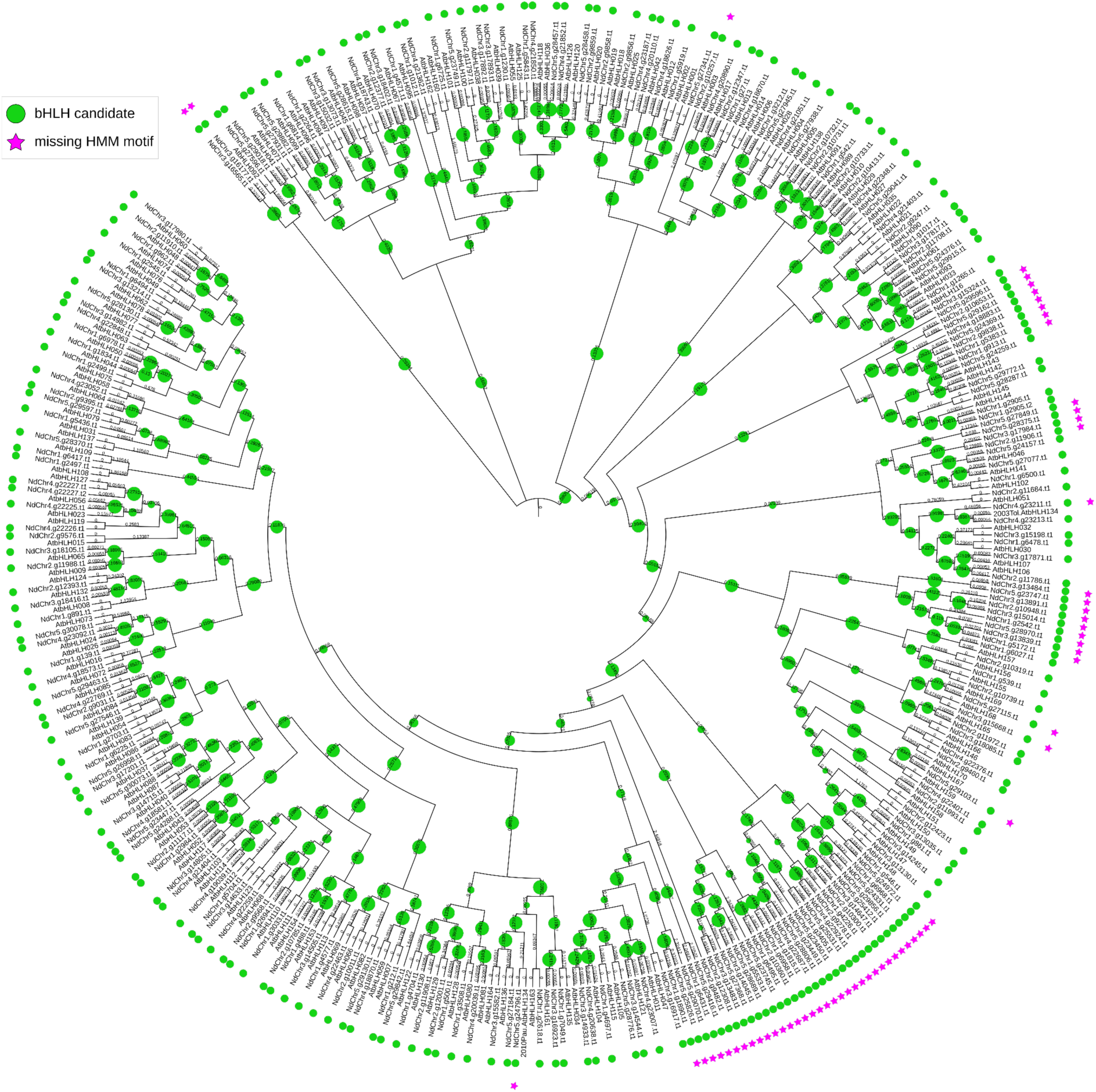
Cladogram derived from a maximum likelihood tree showing the phylogenetic relationship between identified candidates in the A. thaliana accession Nd-1 and A. thaliana bHLHs of the bait collection. Bootstrap values are represented by green circle size. Figure created with iTOL [94]

Of these candidates, 213 were directly assigned to candidates identified in the Col-0 accession. Two other candidates (NdChr1.g2497.t1, NdChr5.g28287.t1) were assigned to the *A. thaliana* bHLHs of the bait collection not identified in Col-0 (AT1G25310 and AT5G50010). The 20 remaining candidates were not assigned to Col-0 orthologs. 18 of these candidates fitted the description of candidates with unclear phylogenetic placement (type 4 candidates in *A. thaliana* Col-0), as they were outliers on very long branches or in groups located on long branches with not more than one bait sequence. Of the identified Col-0 candidates, 30 candidates did not have an assigned Nd-1 candidate. Of these candidates, 27 were outliers with unclear phylogenetic placement (type 4), while one bHLH candidate described in the literature [1, 9, 11] (AT4G28790) and two candidates grouping into bHLH clades (type 3) were without assignment.

#### Monocot species *D. dumetorum*

To test the performance on a crop with huge phylogenetic distance to the model organism *A. thaliana*, the monocotyledonous species *D. dumetorum* was analysed. The pipeline identified 747 initial candidates in *D. dumetorum* [87]. After the first classification, this number was reduced to 254 candidates. The second classification identified 235 possible bHLH candidates (Figure 8). Of these candidates, 200 contained the bHLH motif of the optimised bait collection.

**Figure 8:**
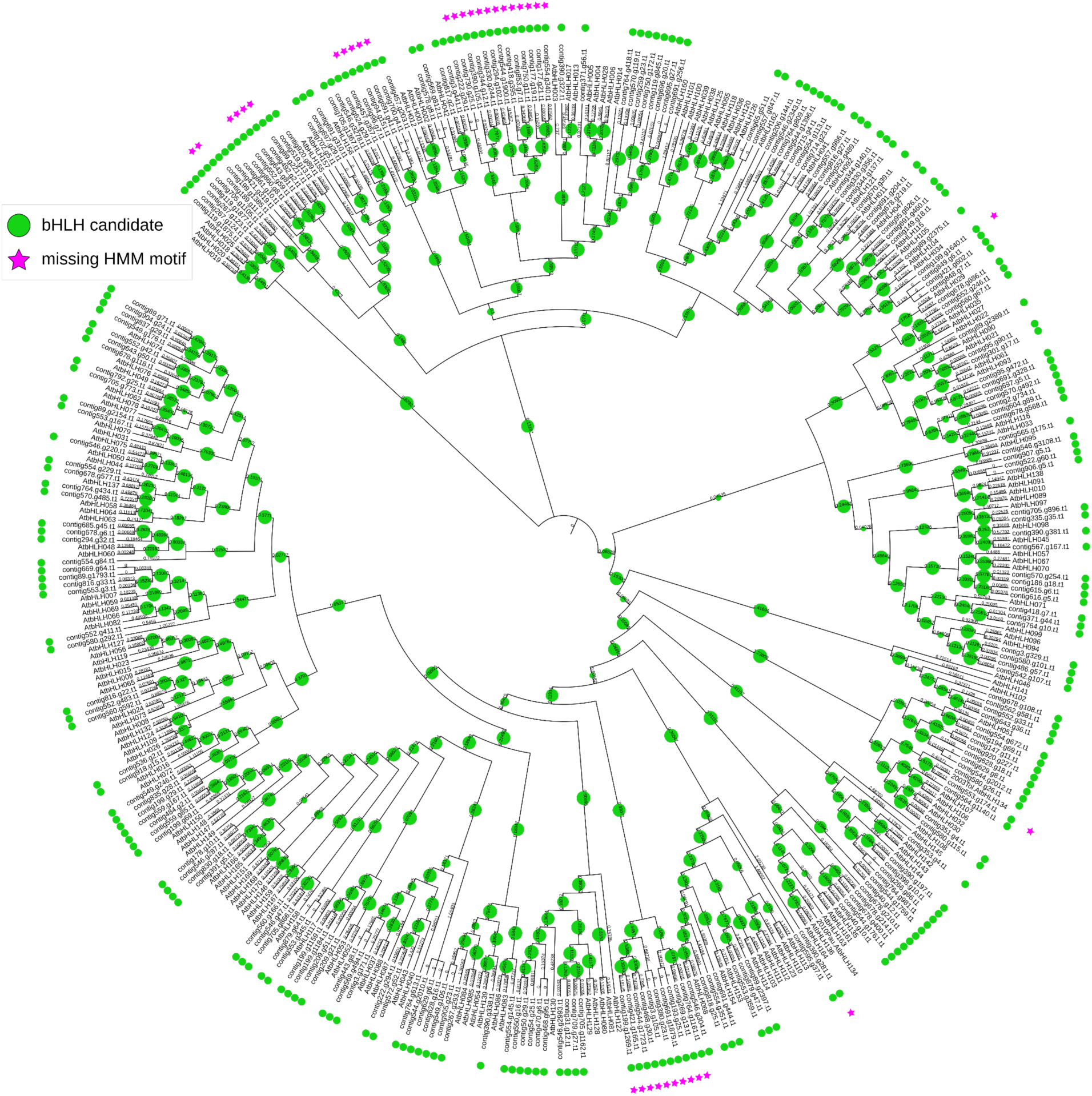
Cladogram derived from a maximum likelihood tree showing the phylogenetic relationship between identified candidates in D. dumetorum and A. thaliana bHLHs of the bait collection. Bootstrap values are represented by green circle size. Figure created with iTOL [94].

#### Transcriptome assembly of *C. tiglium*

A transcriptome assembly was screened for bHLHs to validate the performance of the pipeline on a highly redundant sequence data set. The pipeline identified 1458 initial candidates in the transcriptome assembly of *C. tiglium* [88]. In a first attempt without the parallel option, the pipeline was aborted in the first classification as the alignment with Muscle5 [62] was too memory consuming when aligning the 1893 sequences used for the classification. With the parallel option, the first classification was distributed onto five phylogenetic trees, which reduced the number of candidate sequences per analysis. As a result, 648 candidates were identified. The second classification was distributed onto three phylogenetic trees and identifies 552 possible bHLH candidates of whom 308 harboured the bHLH motif of the bait collection. By utilising the collapsing option of the pipeline, 178 representative candidates for the isoforms of the transcriptome assembly were determined, of whom 120 contain the bHLH motif (Figure 9).

**Figure 9:**
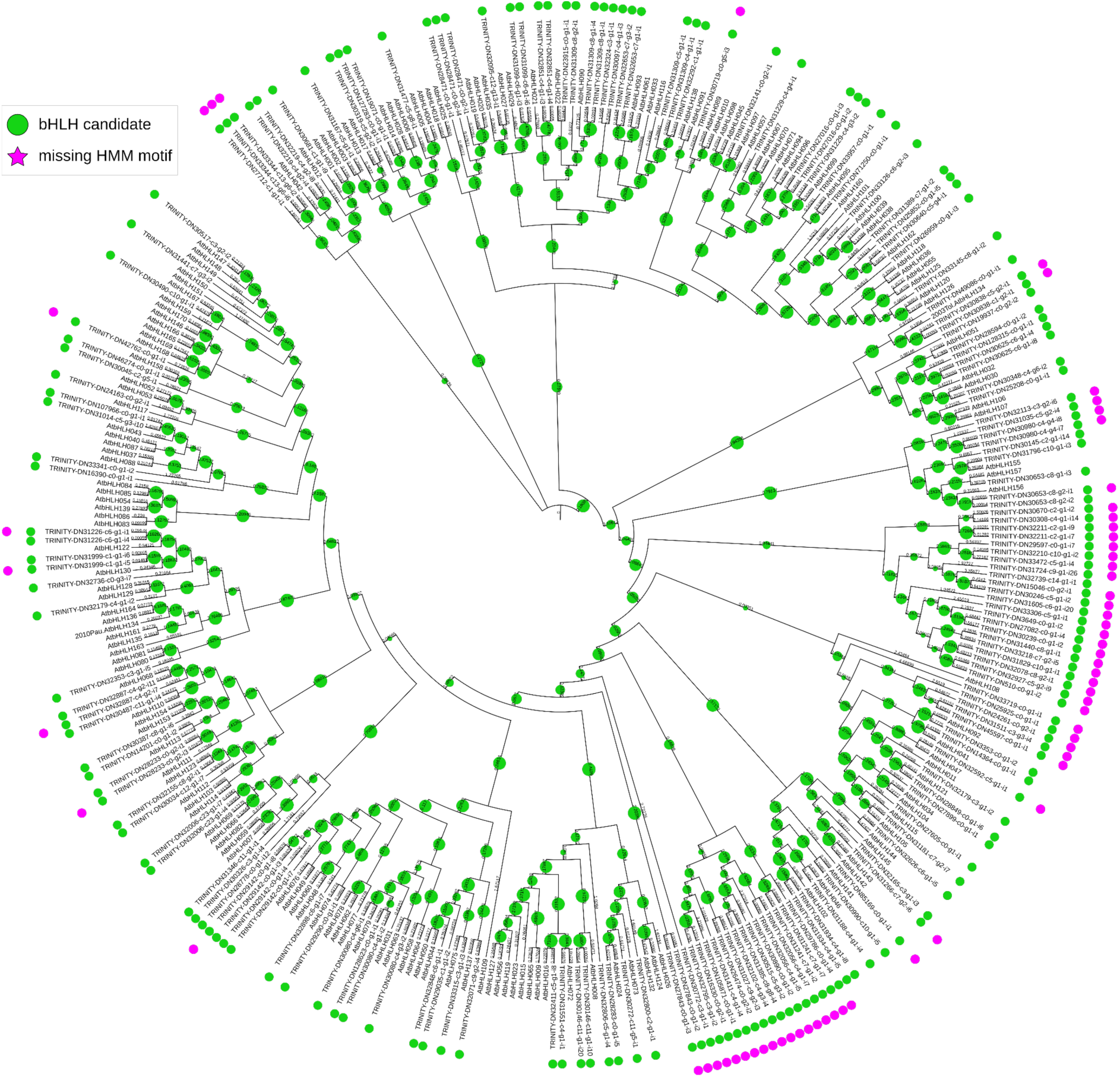
Cladogram derived from a maximum likelihood tree showing the phylogenetic relationship between identified representative candidates in C. tiglium and A. thaliana bHLHs of the bait collection. Bootstrap values are represented by green circle size. Figure created with iTOL [94].

## Discussion

Our bHLH_annotator pipeline enables the automatic identification and annotation of the bHLH gene family in a plant species. The bHLH bait collection covers a wide phylogenetic range of land plant species, including early branching species like *Physcomitrella patens.* As observed in other multi-species studies [10, 11], bHLH sequences from various species cluster together in highly conserved subfamilies. The full diversity of these subfamilies is represented by the optimised bHLH bait collection with a minimal number of sequences. The optimization reduced the computation costs and run time of the analysis substantially. In agreement with the assumption that the bHLH family is monophyletic [9, 10], the outgroup sequences form a distinct clade separated from the bHLH sequences.

A major challenge has been the construction of an accurate multiple sequence alignment due to low sequence conservation in certain regions of bHLHs. Alignments produced by Muscle5 have been shown to lead to phylogenetic trees with the most consistent separation of the outgroup sequences. This is in agreement with the Muscle5 publication that claims a higher accuracy than other established alignment tools [62]. However, Muscle5 has been shown to be time- and resource consuming which makes it less well suited for large datasets [62]. Thus, the number of sequences aligned must be minimised by utilising the optimised bait collection as a minimal, but diversity representing collection of bait sequences for the alignments. The number of initially identified candidates is another factor that contributes to the computational costs. The bHLH_annotator comes with a parallel option, which assigns the initial candidates to separate bins to perform several parallel classifications. The successful deployment of the pipeline on the *C. tiglium* sequences with the parallel option demonstrates the ability to functionally annotate the bHLH transcription factor family in transcriptome assemblies with large numbers of transcript isoforms.

Deploying the pipeline on the *A. thaliana* accession Col-0 resulted in the identification of 165 out of 167 bHLHs. This means that some sequences are missed via BLAST [95, 96]. Because the bHLH family in *A. thaliana* is well studied [1, 9, 11], it is very likely that also bHLHs with a high degree of specialisation and corresponding sequence divergence are described in literature. These sequences can be challenging to identify via similarity to bait sequences due to their high degree of sequence divergence. The addition of diverged *B. napus* and *Z. mays* sequences to the bait collection promised a more extensive identification in closely related monocot and eudicot species. However, HMMER [66] is a sensitive approach to identify even diverged candidate sequences in all species and outperformed BLAST in this respect. Regardless, a comprehensive identification of the bHLH family is demonstrated in the Col-0 accession of *A. thaliana,* which is also supported by the identification of eight additional bHLHs not described in the literature (type 1+2, see results for details). The observation that all but one candidate of the bHLH candidates described in the literature [1, 9, 11] and identified in Col-0 were also identified in Nd-1 demonstrates that the identification is not prevented by a small number of sequence variations.

Through BLAST [95, 96], 68 additional candidates were identified in the Col-0 accession that miss the bHLH domain and are not described in literature. While 13 of these candidates were placed into clades of bHLHs (type 3, see results for details), 55 candidates represented outliers on long branches or were placed into groups of long branches with not more than one bHLH (type 4, see results for details). Some of these were recognized because they were placed between the outgroup sequences and bHLHs in the phylogenetic tree and were not filtered in the classification. There are several explanations for the identification of the remaining candidates. One possible explanation is that members of the bHLH family lost their characteristic domain and perhaps function. A possible example is AT5G50960, which was functionally annotated by TAIR as a nucleotide binding protein that homodimerizes. An argument against the domain loss theory for this protein is the localisation in the cytosol predicted on TAIR [67, 68]. Another explanation is the evolution of the bHLH family via domain shuffling [9–11]. If the affected bHLH gene is the result of domain shuffling, identified candidates can represent orthologs that emerged prior to the insertion of the bHLH encoding exons into the gene. It is also possible that some of the candidates represented pseudogenes, which have also been observed in other studies [11, 31], or artefacts. Furthermore, some candidates can be misplaced in the phylogenetic tree because no member with a close phylogenetic relationship was contained in the bait sequence collection. In total, 30 of the Col-0 bHLH candidates, mainly representing outliers on long branches, were assigned to any identified bHLH candidate in Nd-1. The same applies to 20 candidates in Nd-1. Keeping this observation in mind, the interpretation of candidates on long branches must be performed with high caution and justifies an in-depth investigation of their phylogenetic relationship. The candidates missing the HMM motif and with a phylogenetic placement inside the bHLHs (type 3, see results for details) can be interpreted with a higher certainty. These might represent good candidates for the investigation of domain loss events.

However, if the identification of bHLH candidates with domain loss is not an objective of the study, candidates missing the HMM motif can be excluded by applying the filter domain option or utilising the HMMER [66] search option. As the HMMER [66] search identified all *A. thaliana* bHLHs described in the literature [1, 9, 11] and the additional identified bHLHs (type 1+2, see results for details) in the Col-0 accession, it demonstrated a sensitive identification of sequences harbouring the bHLH domain. This included even diverged sequences and reduced the number of additional candidates.

The bHLH_annotator pipeline was developed to functionally annotate the bHLH transcription factor family. This is also possible with other available tools intended for the functional annotation of sequences: The Pfam database [36, 37] provides the HMM motif PF00010, which represents the bHLH domain [97]. InterProScan5 [97] can be applied to identify members of the bHLHs. Also, motifs like IPR045239 are provided by InterPro, which is described as a transcription factor bHLH95-like bHLH domain, identifying a specific group of plant bHLHs [97]. Although no detailed functional annotation is provided by Pfam, it can be useful for initial identification of bHLHs comparable to the initial results of the bHLH_annotator pipeline. A functional annotation can be assigned to sequences by the Mercator4 pipeline [43, 46]. Sequences assigned to bin 15.5.32 are annotated as bHLHs with no further functional description. An example for a bin providing a functional description is 26.1.2.1.3, which describes bHLHs involved in the regulation of blue light perception, like cryptochrome interacting bHLHs (CIBs) [43, 46]. Also, loss of function can be detected by identifying bins with no assigned sequences [43]. But as the pipeline only provides functional annotation and does not put the sequences into phylogenetic context, no further information can be obtained. A phylogenetic approach is recommended for the annotation of entire gene families [52, 98]. Through the investigation of the phylogenetic relationship, gene duplications can be recognized and functional annotations can be placed into context [52]. Moreover, bHLH family members that experienced loss of function can be identified through a phylogenetic approach. But there are also some pitfalls that must be kept in mind. Functional diversification might cause orthologs to have (slightly) different functions in different species. Furthermore, paralogs can appear as orthologs, if their respective ortholog is lost [49]. OrthoFinder is a tool that enables the phylogenetic identification of orthologs if clear one-to-one relationships exist between genes in different species [54]. But as no direct functional annotation is provided, this step has to be done manually for example by inferring the Gene Ontology (GO) terms of identified orthologs [39, 40].

The bHLH_annotator pipeline was developed for the functional annotations of the bHLH transcription factor family in plants. During conceptualization and implementation, bHLH-specific characteristics have been considered. Additional functionalities are provided by the recognition of subfamily specific motifs and the prediction of DNA binding properties. This allows an automatic and detailed investigation of the bHLH transcription factor family in a wide range of plant species. For future improvement of the pipeline, a more diverse outgroup could possibly lead to a refinement of the candidate classification. Adding reference sequences of other species than *A. thaliana* to the reference collection used for functional annotation would be helpful for the annotation of species that are distantly related to *A. thaliana*. Therefore, users are enabled to add their own collection of landmark bHLH sequences to the set of reference sequences.

## Conclusion

With the bHLH_annotator, a pipeline for the automatic identification and functional annotation of the basic helix-loop-helix (bHLH) transcription factor family in plants is available. Along with the pipeline, a comprehensive collection of bHLH sequences is provided, which represents the full diversity of the bHLH gene family. An optimised collection of the bHLH sequences containing only representative sequences of each bHLH lineage saves time and resources during analysis. The pipeline performs functional annotation through the phylogenetic identification of orthologs, which are often considered to share a common function. Depending on the utilised search option, former bHLHs which experienced domain loss can be identified via BLAST. A sensitive identification limited to bHLHs harbouring the bHLH domain is provided by HMMER. Phylogenetic trees constructed with the optimised bait sequence collection and *A. thaliana* bHLHs allow a detailed investigation of the annotated bHLHs. Characteristics like the subfamily specific motifs and prediction of DNA binding properties are analysed. Furthermore, the bait collections, reference sequences utilised for the annotation, and *A. thaliana* bHLH sequences can be customised for the intended purpose. This provides a powerful set of options for the investigation of the bHLH transcription factor family in land plants, even for transcriptome assemblies. For analysing the evolution of the bHLH family, identifying domain loss events or understanding the development of biological function, a wide range of functions regulated by this transcription factor family can be examined.

## Availability and requirements

Project name: bHLH_annotator

Project home page: https://github.com/bpucker/bHLH_annotor

Operating system(s): Linux

Programming language: Python3

Other requirements: dendropy, BLAST, HMMER, MAFFT, MUSCLE, RAxML or FastTree2

License: GNU General Public License v3.0

Any restrictions to use by non-academics: none

## Declarations

### Ethics approval and consent to participate

Not applicable

### Consent for publication

Not applicable

### Availability of data and materials

The pipeline and the required input data sets are freely available via github: https://github.com/bpucker/bHLH_annotator. Additionally, the released version was archived via zenodo(10.5281/zenodo.7903959). The analysed transcriptomic and genomic resources are publicly available: *Arabidopsis thaliana* Col-0 (https://www.arabidopsis.org/), *Arabidopsis thaliana* Nd-1 (https://doi.org/10.5447/IPK/2019/4), *Dioscorea dumetorum* (https://doi.org/10.4119/unibi/2941469), and *Croton tiglium* (PRJNA416498).

### Authors’ contributions

CT and BP planned the study. CT wrote the software and performed the bioinformatic analysis. CT and BP interpreted the results, and wrote the manuscript. All authors have read the final version of the manuscript and approved its submission.

## Acknowledgements

This work was supported by the BMBF-funded de.NBI Cloud within the German Network for Bioinformatics Infrastructure (de.NBI) (031A532B, 031A533A, 031A533B, 031A534A, 031A535A, 031A537A, 031A537B, 031A537C, 031A537D, 031A538A). Many thanks to the Bioinformatics Resource Facility (BRF) at the Center for Biotechnology (CeBiTec) at Bielefeld University for providing an environment to perform the computational analyses. bioRender.com was used to construct some of the figures.

## References

1. Heim MA, Jakoby M, Werber M, Martin C, Weisshaar B, Bailey PC. The basic helix-loop-helix transcription factor family in plants: a genome-wide study of protein structure and functional diversity. Mol Biol Evol. 2003;20:735–47.

2. Murre C, McCaw PS, Baltimore D. A new DNA binding and dimerization motif in immunoglobulin enhancer binding, daughterless, MyoD, and myc proteins. Cell. 1989;56:777–83.

3. Voronova A, Baltimore D. Mutations that disrupt DNA binding and dimer formation in the E47 helix-loop-helix protein map to distinct domains. Proc Natl Acad Sci U S A. 1990;87:4722–6.

4. Ferré-D’Amaré AR, Prendergast GC, Ziff EB, Burley SK. Recognition by Max of its cognate DNA through a dimeric b/HLH/Z domain. Nature. 1993;363:38–45.

5. Ellenberger T, Fass D, Arnaud M, Harrison SC. Crystal structure of transcription factor E47: E-box recognition by a basic region helix-loop-helix dimer. Genes Dev. 1994;8:970–80.

6. Shimizu T, Toumoto A, Ihara K, Shimizu M, Kyogoku Y, Ogawa N, et al. Crystal structure of PHO4 bHLH domain-DNA complex: flanking base recognition. EMBO J. 1997;16:4689– 97.

7. Murre C, McCaw PS, Vaessin H, Caudy M, Jan LY, Jan YN, et al. Interactions between heterologous helix-loop-helix proteins generate complexes that bind specifically to a common DNA sequence. Cell. 1989;58:537–44.

8. Atchley WR, Fitch WM. A natural classification of the basic helix–loop–helix class of transcription factors. Proc Natl Acad Sci U S A. 1997;94:5172–6.

9. Toledo-Ortiz G, Huq E, Quail PH. The Arabidopsis basic/helix-loop-helix transcription factor family. Plant Cell. 2003;15:1749–70.

10. Pires N, Dolan L. Origin and diversification of basic-helix-loop-helix proteins in plants. Mol Biol Evol. 2010;27:862–74.

11. Carretero-Paulet L, Galstyan A, Roig-Villanova I, Martínez-García JF, Bilbao-Castro JR, Robertson DL. Genome-wide classification and evolutionary analysis of the bHLH family of transcription factors in Arabidopsis, poplar, rice, moss, and algae. Plant Physiol. 2010;153:1398–412.

12. Zhang X-Y, Qiu J-Y, Hui Q-L, Xu Y-Y, He Y-Z, Peng L-Z, et al. Systematic analysis of the basic/helix-loop-helix (bHLH) transcription factor family in pummelo (Citrus grandis) and identification of the key members involved in the response to iron deficiency. BMC Genomics. 2020;21:233.

13. Fairman R, Beran-Steed RK, Anthony-Cahill SJ, Lear JD, Stafford WF, DeGrado WF, et al. Multiple oligomeric states regulate the DNA binding of helix-loop-helix peptides. Proc Natl Acad Sci U S A. 1993;90:10429–33.

14. Wright WE, Catala F, Farmer K. Multimeric structures influence the binding activity of bHLH muscle regulatory factors. Symp Soc Exp Biol. 1992;46:79–87.

15. Baudry A, Heim MA, Dubreucq B, Caboche M, Weisshaar B, Lepiniec L. TT2, TT8, and TTG1 synergistically specify the expression of BANYULS and proanthocyanidin biosynthesis in Arabidopsis thaliana. Plant J Cell Mol Biol. 2004;39:366–80.

16. Albert NW, Davies KM, Lewis DH, Zhang H, Montefiori M, Brendolise C, et al. A Conserved Network of Transcriptional Activators and Repressors Regulates Anthocyanin Pigmentation in Eudicots. Plant Cell. 2014;26:962–80.

17. Albert NW. Subspecialization of R2R3-MYB Repressors for Anthocyanin and Proanthocyanidin Regulation in Forage Legumes. Front Plant Sci. 2015;6.

18. Ramsay NA, Glover BJ. MYB–bHLH–WD40 protein complex and the evolution of cellular diversity. Trends Plant Sci. 2005;10:63–70.

19. Zimmermann IM, Heim MA, Weisshaar B, Uhrig JF. Comprehensive identification of Arabidopsis thaliana MYB transcription factors interacting with R/B-like BHLH proteins. Plant J Cell Mol Biol. 2004;40:22–34.

20. Crooks GE, Hon G, Chandonia J-M, Brenner SE. WebLogo: a sequence logo generator. Genome Res. 2004;14:1188–90.

21. Li X, Huang H, Zhang Z-Q. Genome-wide identification and expression analysis of bHLH transcription factors reveal their putative regulatory effects on petal nectar spur development in Aquilegia. 2022;:2022.04.20.488976.

22. Brownlie P, Ceska T, Lamers M, Romier C, Stier G, Teo H, et al. The crystal structure of an intact human Max-DNA complex: new insights into mechanisms of transcriptional control. Struct Lond Engl 1993. 1997;5:509–20.

23. Atchley WR, Terhalle W, Dress A. Positional dependence, cliques, and predictive motifs in the bHLH protein domain. J Mol Evol. 1999;48:501–16.

24. Winston RL, Gottesfeld JM. Rapid identification of key amino-acid-DNA contacts through combinatorial peptide synthesis. Chem Biol. 2000;7:245–51.

25. Ding A, Ding A, Li P, Wang J, Cheng T, Bao F, et al. Genome-Wide Identification and Low-Temperature Expression Analysis of bHLH Genes in Prunus mume. Front Genet. 2021;12.

26. Fan Y, Yang H, Lai D, He A, Xue G, Feng L, et al. Genome-wide identification and expression analysis of the bHLH transcription factor family and its response to abiotic stress in sorghum [Sorghum bicolor (L.) Moench]. BMC Genomics. 2021;22:415.

27. Bano N, Patel P, Chakrabarty D, Bag SK. Genome-wide identification, phylogeny, and expression analysis of the bHLH gene family in tobacco (Nicotiana tabacum). Physiol Mol Biol Plants Int J Funct Plant Biol. 2021;27:1747–64.

28. Fisher F, Goding CR. Single amino acid substitutions alter helix-loop-helix protein specificity for bases flanking the core CANNTG motif. EMBO J. 1992;11:4103–9.

29. Hyun Y, Lee I. KIDARI, encoding a non-DNA Binding bHLH protein, represses light signal transduction in Arabidopsis thaliana. Plant Mol Biol. 2006;61:283–96.

30. Ledent V, Vervoort M. The basic helix-loop-helix protein family: comparative genomics and phylogenetic analysis. Genome Res. 2001;11:754–70.

31. Li X, Duan X, Jiang H, Sun Y, Tang Y, Yuan Z, et al. Genome-wide analysis of basic/helix-loop-helix transcription factor family in rice and Arabidopsis. Plant Physiol. 2006;141:1167–84.

32. Morgenstern B, Atchley WR. Evolution of bHLH transcription factors: modular evolution by domain shuffling? Mol Biol Evol. 1999;16:1654–63.

33. Bolger ME, Weisshaar B, Scholz U, Stein N, Usadel B, Mayer KFX. Plant genome sequencing - applications for crop improvement. Curr Opin Biotechnol. 2014;26:31–7.

34. Marks RA, Hotaling S, Frandsen PB, VanBuren R. Representation and participation across 20 years of plant genome sequencing. Nat Plants. 2021;7:1571–8.

35. Pucker B, Irisarri I, Vries J de, Xu B. Plant genome sequence assembly in the era of long reads: Progress, challenges and future directions. Quant Plant Biol. 2022;3:e5.

36. Sonnhammer EL, Eddy SR, Durbin R. Pfam: a comprehensive database of protein domain families based on seed alignments. Proteins. 1997;28:405–20.

37. Mistry J, Chuguransky S, Williams L, Qureshi M, Salazar GA, Sonnhammer ELL, et al. Pfam: The protein families database in 2021. Nucleic Acids Res. 2021;49:D412–9.

38. Jones P, Binns D, Chang H-Y, Fraser M, Li W, McAnulla C, et al. InterProScan 5: genome-scale protein function classification. Bioinformatics. 2014;30:1236–40.

39. Ashburner M, Ball CA, Blake JA, Botstein D, Butler H, Cherry JM, et al. Gene Ontology: tool for the unification of biology. Nat Genet. 2000;25:25–9.

40. Gene Ontology Consortium. The Gene Ontology resource: enriching a GOld mine. Nucleic Acids Res. 2021;49:D325–34.

41. Kanehisa M, Furumichi M, Sato Y, Kawashima M, Ishiguro-Watanabe M. KEGG for taxonomy-based analysis of pathways and genomes. Nucleic Acids Res. 2023;51:D587–92.

42. Thimm O, Bläsing O, Gibon Y, Nagel A, Meyer S, Krüger P, et al. MAPMAN: a user-driven tool to display genomics data sets onto diagrams of metabolic pathways and other biological processes. Plant J Cell Mol Biol. 2004;37:914–39.

43. Schwacke R, Ponce-Soto GY, Krause K, Bolger AM, Arsova B, Hallab A, et al. MapMan4: A Refined Protein Classification and Annotation Framework Applicable to Multi-Omics Data Analysis. Mol Plant. 2019;12:879–92.

44. Götz S, García-Gómez JM, Terol J, Williams TD, Nagaraj SH, Nueda MJ, et al. High-throughput functional annotation and data mining with the Blast2GO suite. Nucleic Acids Res. 2008;36:3420–35.

45. Moriya Y, Itoh M, Okuda S, Yoshizawa AC, Kanehisa M. KAAS: an automatic genome annotation and pathway reconstruction server. Nucleic Acids Res. 2007;35 Web Server issue:W182–5.

46. Lohse M, Nagel A, Herter T, May P, Schroda M, Zrenner R, et al. Mercator: a fast and simple web server for genome scale functional annotation of plant sequence data. Plant Cell Environ. 2014;37:1250–8.

47. Bolger ME, Arsova B, Usadel B. Plant genome and transcriptome annotations: from misconceptions to simple solutions. Brief Bioinform. 2018;19:437–49.

48. The UniProt Consortium. UniProt: the universal protein knowledgebase in 2021. Nucleic Acids Res. 2021;49:D480–9.

49. Koonin EV. Orthologs, paralogs, and evolutionary genomics. Annu Rev Genet. 2005;39:309–38.

50. Li L, Stoeckert CJ, Roos DS. OrthoMCL: identification of ortholog groups for eukaryotic genomes. Genome Res. 2003;13:2178–89.

51. Chen F, Mackey AJ, Stoeckert CJ, Roos DS. OrthoMCL-DB: querying a comprehensive multi-species collection of ortholog groups. Nucleic Acids Res. 2006;34 Database issue:D363–368.

52. Eisen JA. Phylogenomics: Improving Functional Predictions for Uncharacterized Genes by Evolutionary Analysis. Genome Res. 1998;8:163–7.

53. Yang Y, Moore MJ, Brockington SF, Soltis DE, Wong GK-S, Carpenter EJ, et al. Dissecting Molecular Evolution in the Highly Diverse Plant Clade Caryophyllales Using Transcriptome Sequencing. Mol Biol Evol. 2015;32:2001–14.

54. Emms DM, Kelly S. OrthoFinder: phylogenetic orthology inference for comparative genomics. Genome Biol. 2019;20:238.

55. Pucker B, Reiher F, Schilbert HM. Automatic Identification of Players in the Flavonoid Biosynthesis with Application on the Biomedicinal Plant Croton tiglium. Plants. 2020;9:1103.

56. Rempel A, Choudhary N, Pucker B. KIPEs3: Automatic annotation of biosynthesis pathways. 2023;:2022.06.30.498365.

57. Emms DM, Kelly S. SHOOT: phylogenetic gene search and ortholog inference. Genome Biol. 2022;23:85.

58. Pucker B. Automatic identification and annotation of MYB gene family members in plants. BMC Genomics. 2022;23:220.

59. Van Rossum G, Drake FL. Python 3 Reference Manual. Scotts Valley, CA: CreateSpace; 2009.

60. Camacho C, Coulouris G, Avagyan V, Ma N, Papadopoulos J, Bealer K, et al. BLAST+: architecture and applications. BMC Bioinformatics. 2009;10:421.

61. Katoh K, Standley DM. MAFFT Multiple Sequence Alignment Software Version 7: Improvements in Performance and Usability. Mol Biol Evol. 2013;30:772–80.

62. Edgar RC. Muscle5: High-accuracy alignment ensembles enable unbiased assessments of sequence homology and phylogeny. Nat Commun. 2022;13:6968.

63. Price MN, Dehal PS, Arkin AP. FastTree 2 – Approximately Maximum-Likelihood Trees for Large Alignments. PLOS ONE. 2010;5:e9490.

64. Kozlov AM, Darriba D, Flouri T, Morel B, Stamatakis A. RAxML-NG: a fast, scalable and user-friendly tool for maximum likelihood phylogenetic inference. Bioinformatics. 2019;35:4453–5.

65. Sukumaran J, Holder MT. DendroPy: a Python library for phylogenetic computing. Bioinforma Oxf Engl. 2010;26:1569–71.

66. Mistry J, Finn RD, Eddy SR, Bateman A, Punta M. Challenges in homology search: HMMER3 and convergent evolution of coiled-coil regions. Nucleic Acids Res. 2013;41:e121.

67. Lamesch P, Berardini TZ, Li D, Swarbreck D, Wilks C, Sasidharan R, et al. The Arabidopsis Information Resource (TAIR): improved gene annotation and new tools. Nucleic Acids Res. 2012;40 Database issue:D1202–10.

68. Cheng C-Y, Krishnakumar V, Chan AP, Thibaud-Nissen F, Schobel S, Town CD. Araport11: a complete reannotation of the Arabidopsis thaliana reference genome. Plant J. 2017;89:789–804.

69. Pucker B, Iorizzo M. Apiaceae FNS I originated from F3H through tandem gene duplication. PLOS ONE. 2023;18:e0280155.

70. Goodstein DM, Shu S, Howson R, Neupane R, Hayes RD, Fazo J, et al. Phytozome: a comparative platform for green plant genomics. Nucleic Acids Res. 2012;40 Database issue:D1178–86.

71. Filiault DL, Ballerini ES, Mandáková T, Aköz G, Derieg NJ, Schmutz J, et al. The Aquilegia genome provides insight into adaptive radiation and reveals an extraordinarily polymorphic chromosome with a unique history. eLife. 2018;7:e36426.

72. Chalhoub B, Denoeud F, Liu S, Parkin IAP, Tang H, Wang X, et al. Plant genetics. Early allopolyploid evolution in the post-Neolithic Brassica napus oilseed genome. Science. 2014;345:950–3.

73. Chen H, Wang T, He X, Cai X, Lin R, Liang J, et al. BRAD V3.0: an upgraded Brassicaceae database. Nucleic Acids Res. 2022;50:D1432–41.

74. Xia E-H, Li F-D, Tong W, Li P-H, Wu Q, Zhao H-J, et al. Tea Plant Information Archive: a comprehensive genomics and bioinformatics platform for tea plant. Plant Biotechnol J. 2019;17:1938–53.

75. Wu GA, Prochnik S, Jenkins J, Salse J, Hellsten U, Murat F, et al. Sequencing of diverse mandarin, pummelo and orange genomes reveals complex history of admixture during citrus domestication. Nat Biotechnol. 2014;32:656–62.

76. Garcia-Mas J, Benjak A, Sanseverino W, Bourgeois M, Mir G, González VM, et al. The genome of melon (Cucumis melo L.). Proc Natl Acad Sci. 2012;109:11872–7.

77. Myburg AA, Grattapaglia D, Tuskan GA, Hellsten U, Hayes RD, Grimwood J, et al. The genome of Eucalyptus grandis. Nature. 2014;510:356–62.

78. Zhang T, Hu Y, Jiang W, Fang L, Guan X, Chen J, et al. Sequencing of allotetraploid cotton (Gossypium hirsutum L. acc. TM-1) provides a resource for fiber improvement. Nat Biotechnol. 2015;33:531–7.

79. Yu J, Jung S, Cheng C-H, Lee T, Zheng P, Buble K, et al. CottonGen: The Community Database for Cotton Genomics, Genetics, and Breeding Research. Plants Basel Switz. 2021;10:2805.

80. Belser C, Baurens F-C, Noel B, Martin G, Cruaud C, Istace B, et al. Telomere-to-telomere gapless chromosomes of banana using nanopore sequencing. Commun Biol. 2021;4:1–12.

81. Edwards KD, Fernandez-Pozo N, Drake-Stowe K, Humphry M, Evans AD, Bombarely A, et al. A reference genome for Nicotiana tabacum enables map-based cloning of homeologous loci implicated in nitrogen utilization efficiency. BMC Genomics. 2017;18:448.

82. Kawahara Y, de la Bastide M, Hamilton JP, Kanamori H, McCombie WR, Ouyang S, et al. Improvement of the Oryza sativa Nipponbare reference genome using next generation sequence and optical map data. Rice. 2013;6:4.

83. Verde I, Abbott AG, Scalabrin S, Jung S, Shu S, Marroni F, et al. The high-quality draft genome of peach (Prunus persica) identifies unique patterns of genetic diversity, domestication and genome evolution. Nat Genet. 2013;45:487–94.

84. Jaillon O, Aury J-M, Noel B, Policriti A, Clepet C, Casagrande A, et al. The grapevine genome sequence suggests ancestral hexaploidization in major angiosperm phyla. Nature. 2007;449:463–7.

85. Woodhouse MR, Cannon EK, Portwood JL, Harper LC, Gardiner JM, Schaeffer ML, et al. A pan-genomic approach to genome databases using maize as a model system. BMC Plant Biol. 2021;21:385.

86. Pucker B, Holtgräwe D, Stadermann KB, Frey K, Huettel B, Reinhardt R, et al. A chromosome-level sequence assembly reveals the structure of the Arabidopsis thaliana Nd-1 genome and its gene set. PLOS ONE. 2019;14:e0216233.

87. Siadjeu C, Pucker B, Viehöver P, Albach DC, Weisshaar B. High Contiguity de novo Genome Sequence Assembly of Trifoliate Yam (Dioscorea dumetorum) Using Long Read Sequencing. Genes. 2020;11:274.

88. Haak M, Vinke S, Keller W, Droste J, Rückert C, Kalinowski J, et al. High Quality de Novo Transcriptome Assembly of Croton tiglium. Front Mol Biosci. 2018;5.

89. Pucker. bHLH annotator. Accessed on: 19.12.2022. https://github.com/bpucker/bHLH_annotator. 2022. https://github.com/bpucker/bHLH_annotator. Accessed 19 Dec 2022.

90. WFO. World Flora Online. 2022. http://www.worldfloraonline.org/. Accessed 4 Apr 2023.

91. The Angiosperm Phylogeny Group, Chase MW, Christenhusz MJM, Fay MF, Byng JW, Judd WS, et al. An update of the Angiosperm Phylogeny Group classification for the orders and families of flowering plants: APG IV. Bot J Linn Soc. 2016;181:1–20.

92. Liu G-Q, Lian L, Wang W. The Molecular Phylogeny of Land Plants: Progress and Future Prospects. Diversity. 2022;14:782.

93. Minh BQ, Schmidt HA, Chernomor O, Schrempf D, Woodhams MD, von Haeseler A, et al. IQ-TREE 2: New Models and Efficient Methods for Phylogenetic Inference in the Genomic Era. Mol Biol Evol. 2020;37:1530–4.

94. Letunic I, Bork P. Interactive Tree Of Life (iTOL) v5: an online tool for phylogenetic tree display and annotation. Nucleic Acids Res. 2021;49:W293–6.

95. Altschul SF, Gish W, Miller W, Myers EW, Lipman DJ. Basic local alignment search tool. J Mol Biol. 1990;215:403–10.

96. Altschul SF, Madden TL, Schäffer AA, Zhang J, Zhang Z, Miller W, et al. Gapped BLAST and PSI-BLAST: a new generation of protein database search programs. Nucleic Acids Res. 1997;25:3389–402.

97. Paysan-Lafosse T, Blum M, Chuguransky S, Grego T, Pinto BL, Salazar GA, et al. InterPro in 2022. Nucleic Acids Res. 2023;51:D418–27.

98. Eisen JA, Wu M. Phylogenetic analysis and gene functional predictions: phylogenomics in action. Theor Popul Biol. 2002;61:481–7.

