## Additional file 1 for "Automatic annotation of the bHLH gene family in plants"

### Identification of highly diverged bHLH sequences

As the bHLH transcription family is investigated in several studies in *Arabidopsis thaliana* [1–3] and *Oryza sativa* [3, 4], it is likely that atypical bHLHs or sequences containing highly diverged domains are identified. To identify these sequences, a sequence similarity search of the bait collection against the polypeptide sequences of the respective species was performed with BLASTp v2.12.0+ [5]. Beforehand, the sequences of the respective species were temporarily removed from the bait collection. Otherwise, they would identify themselves as the best match. The sequences that were not identified via BLAST but are described as bHLH in the literature were specified as the highly diverged sequences of the species. In *A. thaliana*, 10 sequences were specified (AT1G25310.1, AT5G64340.1, AT5G09460.1, AT5G50010.1, AT2G47270.1, AT2G42870.1, AT3G58850.1, AT3G29370.1, AT5G39240.1, AT2G18969.1). In *O. sativa*, 11 sequences were specified (Os01g65080, Os04g35000, Os02g34370, Os02g08220, Os06g44320, Os05g06520, Os02g54870, Os01g43950, Os08g16030, Os04g56500, Os08g31950).

The BLASTp-based Python script “collect_best_BLAST_hits.py” in version v0.29 [6] was used to find the best BLAST hits among the predicted polypeptide sequences of *Brassica napus* for *A. thaliana* bait sequences (GSBRNA2T00086223001,GSBRNA2T00084749001, GSBRNA2T00016048001, GSBRNA2T00101614001, GSBRNA2T00023342001, GSBRNA2T00044957001, GSBRNA2T00114367001, GSBRNA2T00019305001, GSBRNA2T00149121001) and in *Zea mays* for *O. sativa* bait sequences (GRMZM2G047124_P01, GRMZM2G147685_P01, GRMZM2G435001_P01, GRMZM2G364528_P01, GRMZM2G388823_P01, GRMZM6G998221_P01, GRMZM2G075956_P01, GRMZM2G040364_P01, GRMZM2G145909_P01).

### Initial search and classification parameter optimisation

Parameter optimisation was performed based on the *Arabidopsis thaliana* and *Oryza sativa* data sets to avoid overfitting the parameters to one specific species. Before optimising the parameters for a given species, the bait sequences of the respective species were temporarily removed from the bait sequence set. The modified bait and outgroup collections were used as query in a BLASTp v2.12.0+ [5] search with an e-value cutoff of 0.001 against all predicted polypeptide sequences of the respective species. Resulting hits were classified as bHLH and non-bHLH hits, respectively. This was based on whether the sequence identifiers of BLAST hits were contained in the bHLH annotation of the respective species. Both, the bHLH and non-bHLH hit groups, were filtered independently on bit score, alignment length and percentage of identical matches (described as similarity). For values ranging from 0 to 100 in steps of 10 for each of these parameters, the number of remaining hits was determined. The parameter with the highest impact on the results was set to a fixed value and the number of remaining hits determined for all possible combinations of the fixed parameter and the ranges of the two flexible parameters. The results were visualized in a three-dimensional diagram (Figure 1) and the parameters reducing the number of false-positive candidates without a significant loss of bHLH candidates were manually identified.

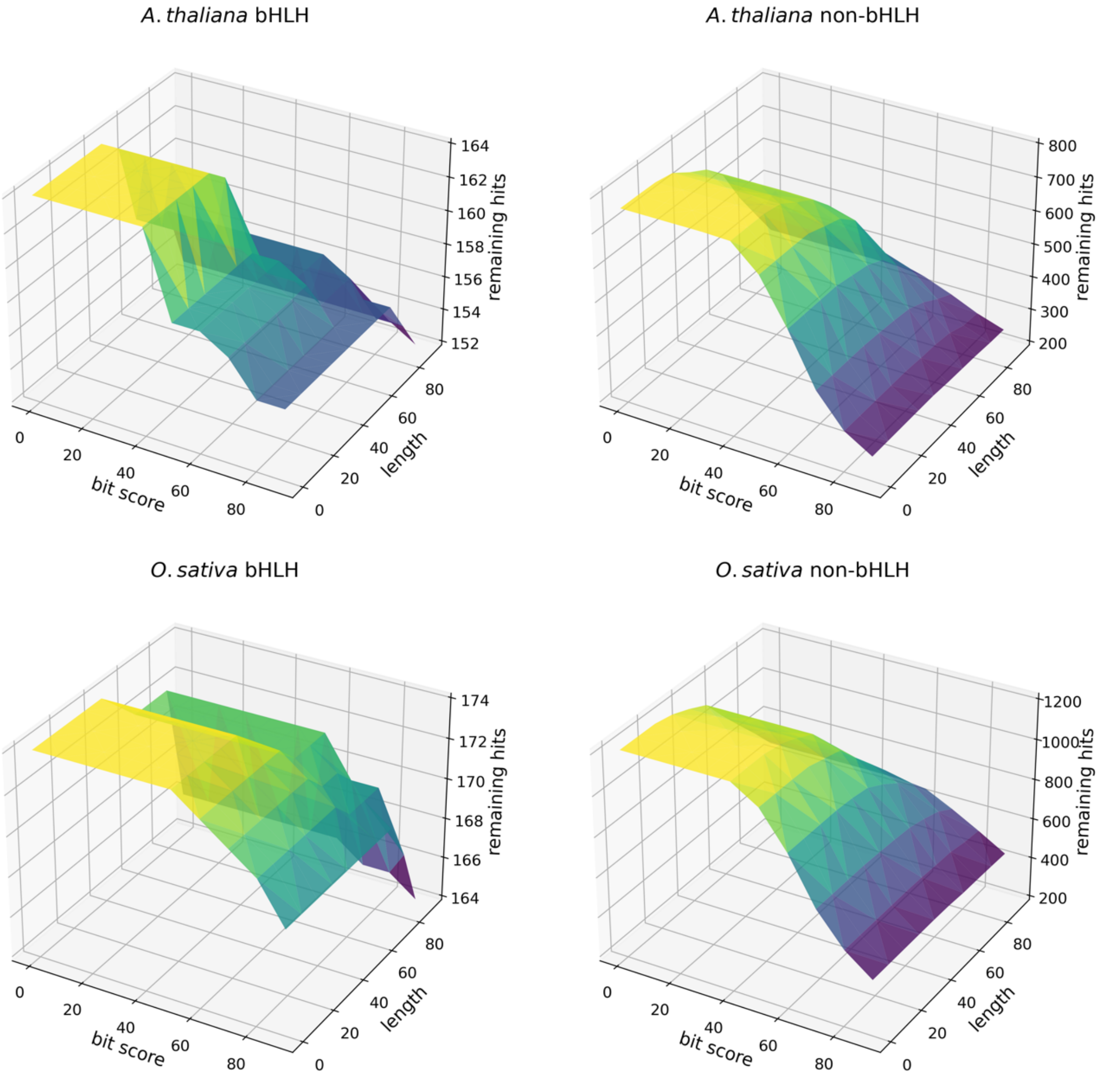


Figure 1: Number of remaining bHLH and non-bHLH hits in *A. thaliana* and *O. sativa* after filtered by varying bit score and length, while using a fixed similarity cutoff of 40.

The optimal parameters for the classification would strongly reduce the number of false-positive candidates while not excluding *bona fide* bHLH candidates. The optimisation of these parameters was performed independently for both species with the already optimised BLAST parameters. The classification (step 2 of the pipeline) was performed while varying the score and the number of neighbouring leaves that are selected for the evaluation of each candidate leaf and identified ingroup candidates classified as bHLHs and non-bHLHs. The results were visualized in a three-dimensional diagram (Figure 2) to identify the optimal parameters that result in best performance on both benchmarking species.

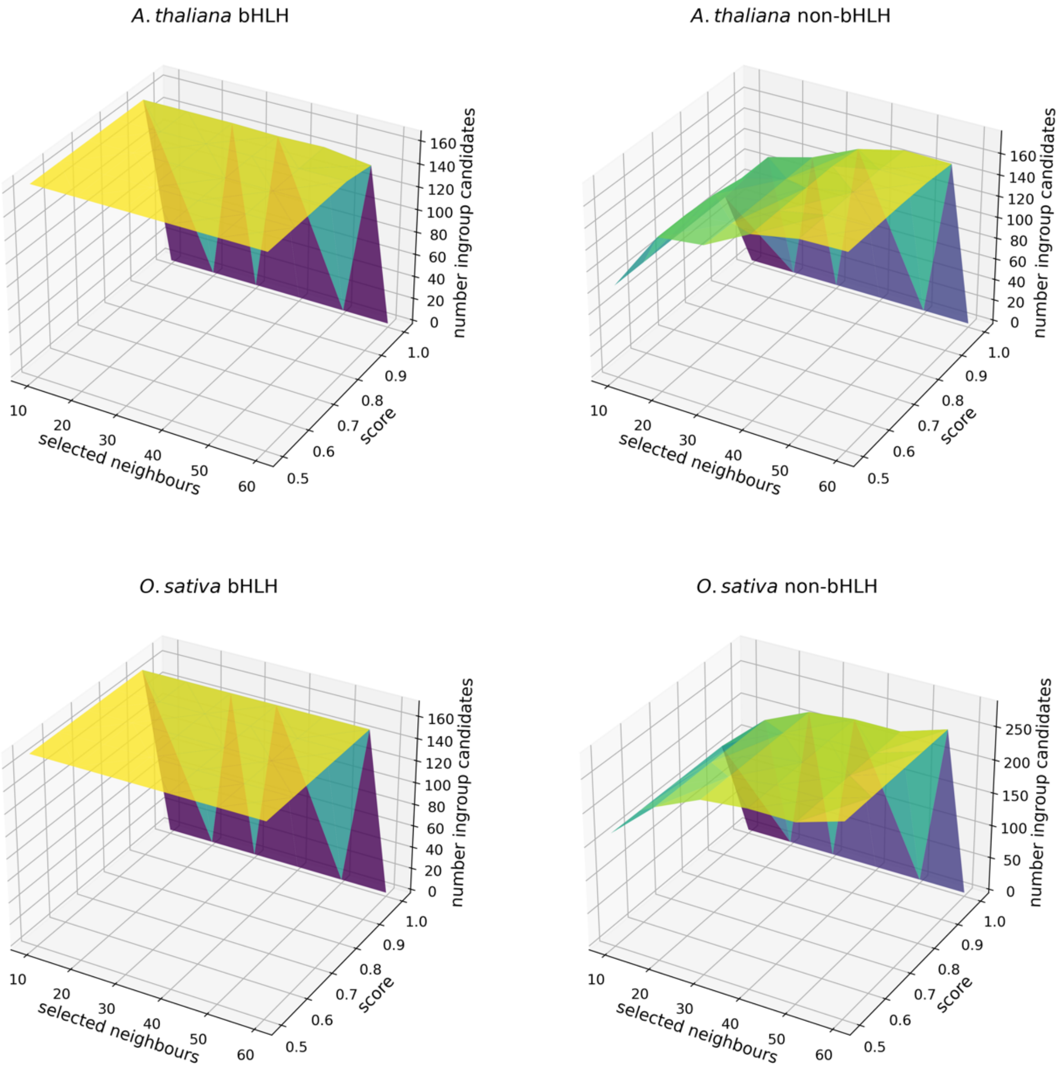


Figure 2: Number of remaining bHLH and non-bHLH ingroup candidates in *A. thaliana* and *O. sativa* for second classification with a varying score and number of selected neighbour leaves.

### References

1. Heim MA, Jakoby M, Werber M, Martin C, Weisshaar B, Bailey PC. The basic helix-loop-helix transcription factor family in plants: a genome-wide study of protein structure and functional diversity. Mol Biol Evol. 2003;20:735–47.

2. Toledo-Ortiz G, Huq E, Quail PH. The Arabidopsis Basic/Helix-Loop-Helix Transcription Factor Family[W]. Plant Cell. 2003;15:1749–70.

3. Carretero-Paulet L, Galstyan A, Roig-Villanova I, Martínez-García JF, Bilbao-Castro JR, Robertson DL. Genome-Wide Classification and Evolutionary Analysis of the bHLH Family of Transcription Factors in Arabidopsis, Poplar, Rice, Moss, and Algae. Plant Physiol. 2010;153:1398–412.

4. Li X, Duan X, Jiang H, Sun Y, Tang Y, Yuan Z, et al. Genome-Wide Analysis of Basic/Helix-Loop-Helix Transcription Factor Family in Rice and Arabidopsis. Plant Physiol. 2006;141:1167–84.

5. Camacho C, Coulouris G, Avagyan V, Ma N, Papadopoulos J, Bealer K, et al. BLAST+: architecture and applications. BMC Bioinformatics. 2009;10:421.

6. Pucker B, Iorizzo M. Apiaceae FNS I originated from F3H through tandem gene duplication. 2022;:2022.02.16.480750.
