## Additional file 2 for "Automatic annotation of the bHLH gene family in plants"

### Identification of outgroup sequences

In order to distinguish between bHLH-like candidates and *bona fide* bHLHs, a collection of phylogenetically close outgroup sequences is needed. Possible outgroup sequences in *A. thaliana* were identified by performing a BLASTp v2.12.0+ [1] search against all predicted polypeptide sequences of TAIR10 using the *A. thaliana* bHLH sequences [2–4] as baits. BLAST hits with a bit score above 50 were considered as bHLH-like candidates. Together with the *A. thaliana* bHLH sequences, the candidates were globally aligned using Muscle5 v5.1.osx64 [5]. The alignment was trimmed by removing positions with less than 10% occupancy at a given alignment position. A maximum likelihood tree was constructed via FastTree v2.1.10 [6] using the "-wag" option. For inspection, the phylogenetic tree was visualised using iTOL [7].

Candidates forming a monophyletic group distinct from the *bona fide* bHLHs were identified as non-bHLH outgroup sequences. These outgroup sequences were identified in all species that are listed in Table 1. For each species, the *A. thaliana* outgroup sequences were included in the BLASTp search in addition to the bHLH sequences of that respective species. BLAST hits revealed outgroup sequences by showing a close phylogenetic relationship to the *A. thaliana* outgroup sequences, but not to the bHLH sequences of the respective species.

Table 1: Plant polypeptide sequence datasets used to collect bHLH sequences. The species, version and database source are given.

| **Species** | **Version** | **Source** |
|---|---|---|
| *Aquilegia coeruela* | v3.1 | Phytozome (Goodstein *et al.*, 2012; Filiault *et al.*, 2018) |
| *Arabidopsis thaliana* | TAIR10 | Phytozome (Lamesch *et al.*, 2012) |
| *Brassica rapa* | v1.2 | Brassicaceae Database (Chen *et al.*, 2022) |
| *Camellia sinesis* | - | Tea Plant Information Archive (Xia *et al.*, 2019) |
| *Citrus grandis* | v1.0 | Phytozome (Goodstein *et al.*, 2012; Wu *et al.*, 2014) |
| *Cucumis melo* | v3.5.1 | Cucurbit Genomics Database (Garcia-Mas *et al.*, 2012) |
| *Eucalyptus grandis* | v2.0 | Phytozome (Goodstein *et al.*, 2012; Myburg *et al.*, 2014) |
| *Gossypium hirsutum* | "TM-1" genome NAU-NBI_v1.1 | Cottongen (Zhang *et al.*, 2015; Yu *et al.*, 2021) |
| *Musa acuminata* | DH-Pahang v4 | Banana Genome Hub (Belser *et al.*, 2021) |
| *Nicotiana tabacum* | v4.5 | SolGenomicsNetwork (Edwards *et al.*, 2017) |
| *Oryza sativa* | 7.0 | Rice Genome Annotation Project (Kawahara *et al.*, 2013) |

| | | |
|---|---|---|
| *Phaseolus vulgaris* | v2.1 | Phytozome (Goodstein *et al.*, 2012) |
| *Physcomitrella patens* | JGI v1.1 | Phytozome (Goodstein *et al.*, 2012) |
| *Populus trichocarpa* | v1.1 | Phytozome (Goodstein *et al.*, 2012) |
| *Prunus persica* | v2.1 | Phytozome (Goodstein *et al.*, 2012; Verde *et al.*, 2013) |
| *Vitis vinifera* | v2.1 | Phytozome (Goodstein *et al.*, 2012; Jaillon *et al.*, 2007) |

### Optimisation of sequence collections through thinning

Large and redundant sequence collections lead to high computational costs and long run times in the following analyses. It is possible to optimise these collections by reducing large groups of very similar sequences to only one representative sequence. The initial bait collection and the initial outgroup collection were separately optimised by thinning based on phylogenetic distance of the individual sequences to generate a small set of sequences that still represent the full phylogenetic diversity of bHLHs. Phylogenetic trees of the collections were constructed with FastTree 2.1.10 [6] as described above. DendroPy 4.5.2 [8] was deployed to calculate the mean nearest taxon distance and patristic distances between all leaves of the trees. For each leaf, neighbouring leaves with a patristic distance less than the mean nearest taxon distance multiplied by a given factor were identified as closely related group members. Leaves identified as group members were excluded from further group member identifications to prevent overlapping. The leaf with the longest sequence was chosen as representative of the group and added to the optimised collection. Leaves with no phylogenetic neighbour in the mean nearest taxon distance multiplied by ten were identified as singular sequences on extraordinarily long branches and excluded from the optimised collection.

The collection optimisation was performed iteratively. Each step included the construction of a phylogenetic tree followed by thinning. First, sequences were thinned per species. Leaves within a patristic distance less than the mean nearest taxon distance multiplied by factor two were considered as a group. Only the longest sequence was retained to represent this group of paralogous sequences in the following steps. The obtained representative sequences from all species were merged and three rounds of thinning were performed. In these steps, the factor for the mean nearest taxon distance was set to one. The representative sequences from the last step were obtained as the optimised collection, a diverse set of sequences with low lineage redundancy that still allows the identification of lineage-specific bHLH sequences. The HMMER 3.3.2 [9] program “hmmbuild” was used to create a HMM motif of the optimised bait collection.

**References**


1. Camacho C, Coulouris G, Avagyan V, Ma N, Papadopoulos J, Bealer K, et al. BLAST+: architecture and applications. BMC Bioinformatics. 2009;10:421.

2. Heim MA, Jakoby M, Werber M, Martin C, Weisshaar B, Bailey PC. The basic helix-loop-helix transcription factor family in plants: a genome-wide study of protein structure and functional diversity. Mol Biol Evol. 2003;20:735–47.

3. Toledo-Ortiz G, Huq E, Quail PH. The Arabidopsis Basic/Helix-Loop-Helix Transcription Factor Family[W]. Plant Cell. 2003;15:1749–70.

4. Carretero-Paulet L, Galstyan A, Roig-Villanova I, Martínez-García JF, Bilbao-Castro JR, Robertson DL. Genome-Wide Classification and Evolutionary Analysis of the bHLH Family of Transcription Factors in Arabidopsis, Poplar, Rice, Moss, and Algae. Plant Physiol. 2010;153:1398–412.

5. Edgar RC. High-accuracy alignment ensembles enable unbiased assessments of sequence homology and phylogeny. 2022;:2021.06.20.449169.

6. Price MN, Dehal PS, Arkin AP. FastTree 2--approximately maximum-likelihood trees for large alignments. PloS One. 2010;5:e9490.

7. Letunic I, Bork P. Interactive Tree Of Life (iTOL) v5: an online tool for phylogenetic tree display and annotation. Nucleic Acids Res. 2021;49:W293–6.

8. Sukumaran J, Holder MT. DendroPy: a Python library for phylogenetic computing. Bioinforma Oxf Engl. 2010;26:1569–71.

9. Mistry J, Finn RD, Eddy SR, Bateman A, Punta M. Challenges in homology search: HMMER3 and convergent evolution of coiled-coil regions. Nucleic Acids Res. 2013;41:e121.
