## Additional file 3 for "Automatic annotation of the bHLH gene family in plants"

### Phylogenetic analysis of bait collection

The bait collection and the optimised bait collection were analysed for the subfamilies described by the previous multi species-study of the bHLH transcription factor family conducted by Pires and Dolan [1]. The bait collection was further analysed for the representation of the lineages Bryophytes, Lycophytes, Gymnosperm, Amborellales, Monocots, and Eudicots in the assigned subfamilies and an Weblogo was created for each subfamily. The assigned subfamilies of the optimised bait collection were compared in detail to the subfamilies described by Pires and Dolan [1] to identify bHLHs which are placed into different subfamilies.

In a first step, the subfamilies were identified in the optimised bait collection. To be able to compare our results with previous multi-species studies of the bHLH transcription factor family [1], the optimised bait collection was extended by all *Arabidopsis thaliana* bHLHs from the bait collection, as some sequences were removed in the optimisation process (see Additional file 2). The resulting 493 sequences were globally aligned using Muscle5 v5.1. [2]. The alignment was trimmed by removing positions with less than 10% occupancy at a given alignment position and a maximum likelihood tree was constructed via FastTree v2.1.10 [3] using the "-wag" option. For inspection, the phylogenetic tree was visualised using iTOL [4]. The subfamilies were identified based on the assigned subfamilies of the *A. thaliana* bHLHs by Pires and Dolan [1].
Because the whole bait collection consists of too many sequences to be aligned by Muscle5 v5.1. [2], the first tree that was received in the optimisation process after the thinning per species was used as representative for the bait collection (see Additional file 2). The subfamilies were assigned for each leaf based on the assigned subfamilies of the optimised bait collection by identifying the next neighbouring leaf of the optimised bait collection with a minimum edge distance and patristic distance in the phylogenetic tree with DendroPy 4.5.2 [5]. The phylogenetic tree was visualised using iTOL [4] and the subfamilies were assigned. To create an Weblogo of each identified subfamily, the subfamily member sequences were globally aligned using Muscle5 v5.1. [2] and positions with less than 80% occupancy were removed from the alignment. The alignment was visualised using Weblogo 3.7.12 [6].

#### Identification of subfamilies in optimised bait collection

In total, 27 subfamilies were identified in the resulting phylogenetic tree (see Additional file 4). All 26 subfamilies described by Pires and Dolan [1] were recovered in our analysis. Also, an additional subfamily formed by previously described orphan sequences (SmbHLH007, AtbHLH021, AtbHLH022, AtbHLH090) and two sequences from the optimised bait collection (BrabHLH125, BrabHLH081) was identified, which is marked as "O" in the phylogenetic tree. The relationship between the subfamilies does not correspond to the analysis of Pires and Dolan [1], which conforms with the findings of other studies that proposed a high conservation for the bHLH subfamilies themself, but not the relationship between them [1, 7–12]. For a detailed investigation, the assigned subfamilies of 218 bHLHs (156 *A. thaliana*, 25 *Physcomitrella patens*, 21 *Selaginella moellendorffii* and 16 *Oryza sativa* bHLHs) included in the analysis performed by Pires and Dolan [1] and the optimised bait collection were compared. Of these sequences, 189 bHLHs are placed into the same subfamily in both analyses. The remaining 29 bHLHs are assigned to different subfamilies, with 20 of them representing previously described orphan sequences (Table 1).

Table 1: bHLHs with differences in the assigned subfamilies between the analysis of the optimised bait collection (this study) and the Pires and Dolan [1] study.

| | **Subfamily in Pires and Dolan [1] study** | **Subfamily in optimised bait collection** |
|---|---|---|
| SmbHLH035 | IVd | Ib(1) |
| SmbHLH036 | IVd | Ib(1) |
| SmbHLH071 | VIIIa | VIIIb |
| SmbHLH092 | IVd | IIIf |
| OsbHLH147 | Ib(2) | IVa |
| PpbHLH055 | VIIIa (named PpbHLH081) | VIIIc(1) |
| PpbHLH059 | VIIIc(2) (named PpbHLH083) | VIIIc(1) |
| PpbHLH076 | XII (named PpbHLH051) | Ia |
| AtbHLH026 | VII(a+b) | XIII |
| AtbHLH108 | Orphans | VII(a+b)(1) |
| AtbHLH109 | Orphans | VII(a+b)(1) |
| AtbHLH147 | Orphans | VIIIa(2) |
| AtbHLH148 | Orphans | VIIIa(2) |
| AtbHLH149 | Orphans | VIIIa(2) |
| AtbHLH150 | Orphans | VIIIa(2) |
| AtbHLH151 | Orphans | VIIIa(2) |
| AtbHLH158 | Orphans | VIIIa(2) |
| AtbHLH159 | Orphans | VIIIa(2) |
| SmbHLH022 | Orphans | VIIIa(2) |
| SmbHLH040 | Orphans | VIIIc(1) |
| SmbHLH070 | Orphans | XV |
| OsbHLH157 | Orphans | VIIIa(2) |
| OsbHLH166 | Orphans | IIIb |
| PpbHLH006 | Orphans (named PpbHLH090) | Ia |
| PpbHLH023 | Orphans (named PpbHLH024) | Ia |
| PpbHLH033 | Orphans (named PpbHLH040) | Ia |
| PpbHLH039 | Orphans (named PpbHLH038) | Ia |
| PpbHLH095 | Orphans (named PpbHLH087) | IVd |
| PpbHLH098 | Orphans (named PpbHLH050) | VIIIc(1) |

### Lineage representation in subfamilies

All 27 subfamilies were recovered in the representative tree of the bait collection. A Weblogo was created for each subfamily, demonstrating the variable position of the bHLH domain between the subfamilies [1, 7, 10, 11] (see Additional file 5). For each bHLH in the phylogenetic tree, the lineage of the species and the assigned subfamily were documented (see Additional file 6).

The representation of the lineages Bryophytes, Lycophytes, Monocots, and Eudicots confirms with the findings of Pires and Dolan [1] except for a missing representation of the Lycophyte lineage in the subfamilies IVb and IVd. For the subfamily IVb, the representative bHLH SmbHLH078 is not included in the phylogenetic tree. For the subfamily IVd, the representative bHLHs SmbHLH020, SmbHLH043, SmbHLH066, and SmbHLH077 are not included in the phylogenetic tree and the representative bHLHs SmbHLH035, SmbHLH036, and SmbHLH092 are not part of the IVd subfamily, as it was observed in the optimised bait collection (Table 1). In addition, the bait collection was also analysed for the representation of the Gymnosperm and Amborellales lineages in the subfamilies.

Corresponding to other multi-species studies [1, 7], most subfamilies are conserved among most of the land plant lineages and all subfamilies are conserved in the angiosperm lineage. In total, the Bryophytes lineage is represented in 20 subfamilies, the Lycophytes and Gymnosperm lineages are represented in 21 subfamilies respectively, and the Amborellales lineage is represented in 18 subfamilies (Table 2).

Table 2: Lineage representation of bHLHs from Bryophytes, Lycophytes, Gymnosperm, Amborellales, Monocots, and Eudicots in the assigned subfamilies of the bait collection.

| | **Bryophytes** | **Lycophytes** | **Gymnosperm** | **Amborellales** | **Monocots** | **Eudicots** |
|---|---|---|---|---|---|---|
| Ia | ● | ● | ● | ● | ● | ● |
| Ib(1) | ● | ● | ● | ● | ● | ● |
| Ib(2) | | | ● | ● | ● | ● |
| II | ● | ● | ● | ● | ● | ● |
| III(a+c) | ● | ● | ● | | ● | ● |
| III(b) | ● | ● | ● | | ● | ● |
| III(d+e) | ● | | ● | ● | ● | ● |
| IIIf | ● | ● | ● | ● | ● | ● |
| IVa | ● | ● | ● | | ● | ● |
| IVb | ● | | ● | ● | ● | ● |
| IVc | ● | ● | | | ● | ● |
| IVd | ● | | ● | ● | ● | ● |
| Va | ● | ● | | ● | ● | ● |
| Vb | | ● | ● | ● | ● | ● |
| VII(a+b) | ● | ● | ● | ● | ● | ● |
| VIII(a) | ● | ● | | ● | ● | ● |
| VIIIb | ● | ● | ● | ● | ● | ● |
| VIIIc(1) | ● | ● | ● | | ● | ● |
| VIIIc(2) | ● | ● | | | ● | ● |
| IX | ● | ● | ● | ● | ● | ● |
| X | | ● | ● | ● | ● | ● |
| XI | ● | ● | ● | ● | ● | ● |
| XII | ● | ● | ● | ● | ● | ● |
| XIII | | ● | | | ● | ● |
| XIV | | ● | ● | ● | ● | ● |

| | | | | | | |
|---|---|---|---|---|---|---|
| XV | | | ● | ● | ● | ● |
| Orphans | ● | ● | | | ● | ● |

For the optimised bait collection, some lineage representing bHLHs were filtered from the subfamilies at the optimisation process due to close phylogenetic relationship to other bHLH sequences. This regards especially bHLHs representing the Gymnosperm, Lycophytes, and Bryophytes, as the bait collection is dominated by bHLHs representing the angiosperm lineage. Keeping this in mind, no conclusions about the lineage representation in the subfamilies can be drawn based on the analysis of the optimised bait collection.

**Conclusion**

Overall, all subfamilies proposed by Pires and Dolan [1] were able to be reconstructed in the phylogenetic analysis of the bait collection. In correspondence to other multi-species studies [1, 7], the subfamilies are highly conserved among the land plant lineages and it was able to confirm the conservation also for the Amborellales and Gymnosperm lineages. The comparation of the subfamily members between Pires and Dolan [1] and the assigned subfamilies in the optimised bait collection demonstrates that the optimisation of the bait collection successfully conserved all important phylogenetic subfamilies of the bHLHs in the thinning process. This supports the application of the optimised bait collection for the automated identification and annotation of plant bHLHs in the bHLH_annotator.

**References**

1. Pires N, Dolan L. Origin and diversification of basic-helix-loop-helix proteins in plants. Mol Biol Evol. 2010;27:862–74.

2. Edgar RC. Muscle5: High-accuracy alignment ensembles enable unbiased assessments of sequence homology and phylogeny. Nat Commun. 2022;13:6968.

3. Price MN, Dehal PS, Arkin AP. FastTree 2--approximately maximum-likelihood trees for large alignments. PloS One. 2010;5:e9490.

4. Letunic I, Bork P. Interactive Tree Of Life (iTOL) v5: an online tool for phylogenetic tree display and annotation. Nucleic Acids Res. 2021;49:W293–6.

5. Sukumaran J, Holder MT. DendroPy: a Python library for phylogenetic computing. Bioinforma Oxf Engl. 2010;26:1569–71.

6. Crooks GE, Hon G, Chandonia J-M, Brenner SE. WebLogo: a sequence logo generator. Genome Res. 2004;14:1188–90.

7. Carretero-Paulet L, Galstyan A, Roig-Villanova I, Martínez-García JF, Bilbao-Castro JR, Robertson DL. Genome-Wide Classification and Evolutionary Analysis of the bHLH Family of Transcription Factors in Arabidopsis, Poplar, Rice, Moss, and Algae. Plant Physiol. 2010;153:1398–412.

8. Atchley WR, Fitch WM. A natural classification of the basic helix–loop–helix class of

transcription factors. Proc Natl Acad Sci. 1997;94:5172–6.

9. Ledent V, Vervoort M. The basic helix-loop-helix protein family: comparative genomics and phylogenetic analysis. Genome Res. 2001;11:754–70.

10. Heim MA, Jakoby M, Werber M, Martin C, Weisshaar B, Bailey PC. The basic helix-loop-helix transcription factor family in plants: a genome-wide study of protein structure and functional diversity. Mol Biol Evol. 2003;20:735–47.

11. Toledo-Ortiz G, Huq E, Quail PH. The Arabidopsis Basic/Helix-Loop-Helix Transcription Factor Family[W]. Plant Cell. 2003;15:1749–70.

12. Li X, Duan X, Jiang H, Sun Y, Tang Y, Yuan Z, et al. Genome-Wide Analysis of Basic/Helix-Loop-Helix Transcription Factor Family in Rice and Arabidopsis. Plant Physiol. 2006;141:1167–84.
