## Additional file 9 for "Automatic annotation of the bHLH gene family in plants"

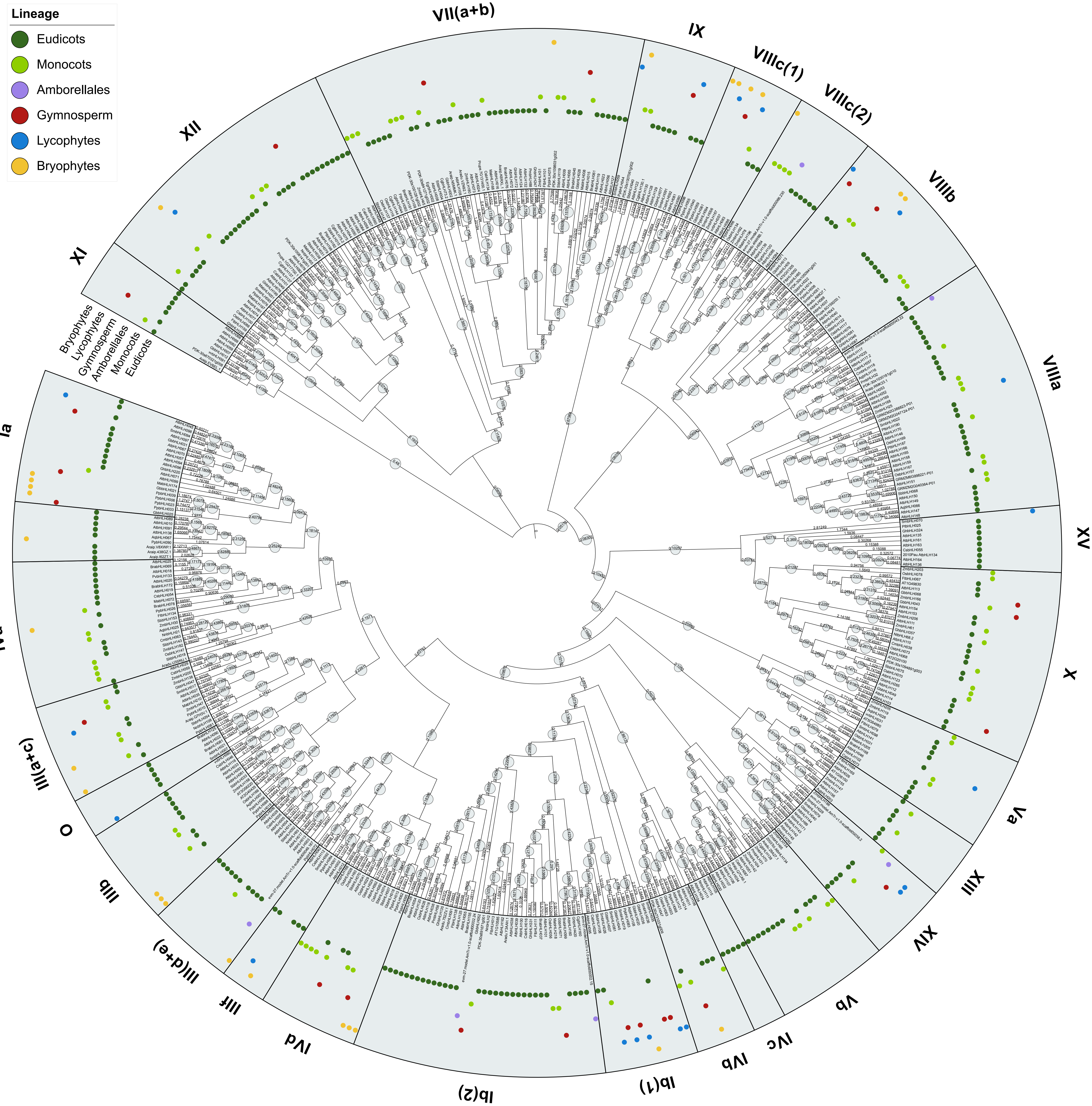


Figure S9: Maximum likelihood tree created with FastTree v2.1.10 showing the phylogenetic relationship between optimised bHLH bait collection v1.1. and *A. thaliana* bHLHs. Twenty-seven bHLH subfamilies were identified and analysed for the included land plant lineages. Branch lengths are displayed. Bootstrap values are represented by the size of the grey circles. Figure created with iTOL.
