## Additional file 10 for "Automatic annotation of the bHLH gene family in plants"

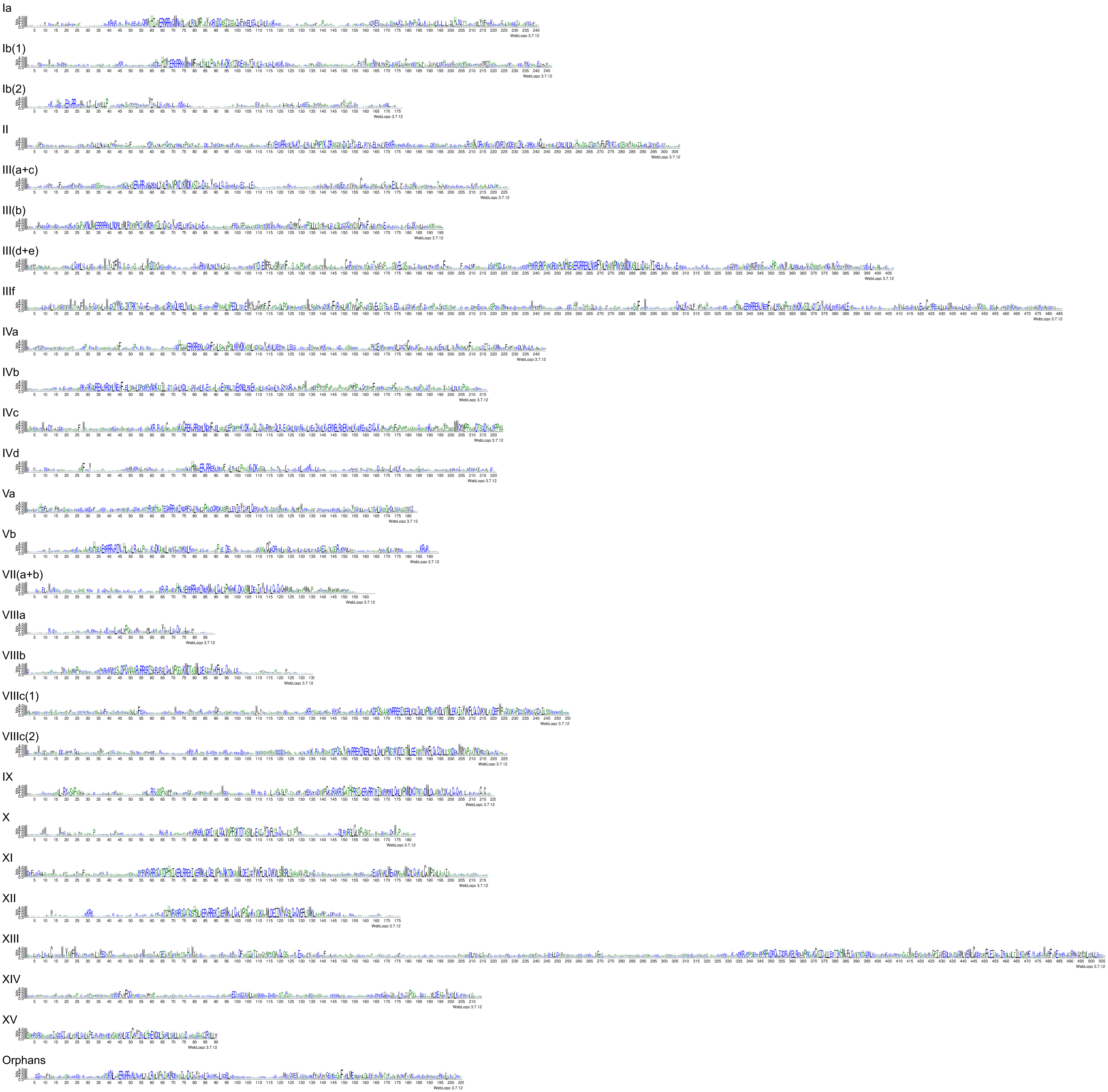


Figure S5: Weblogos of the 27 subfamilies identified in the bait collection. The subfamily member sequences were globally aligned using Muscle5 v5.1. and positions with less than 80% occupancy were removed from the alignment. Weblogos created with WebLogo 3.7.12.
