## Additional file 12 for "Automatic annotation of the bHLH gene family in plants"

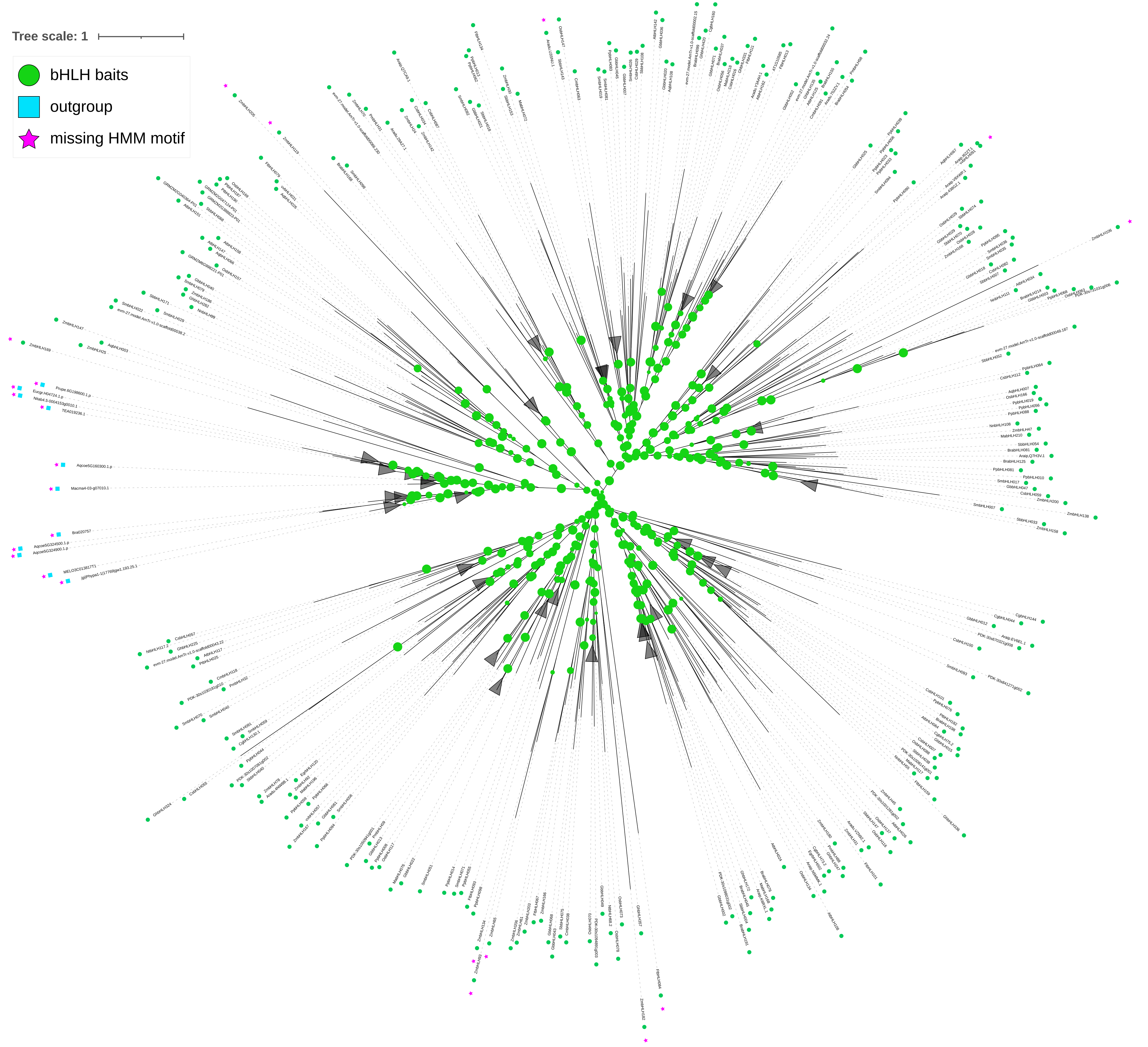


Figure S12: Maximum likelihood tree created with FastTree v2.1.10 showing the phylogenetic relationship between optimised bHLH bait collection and optimised outgroup collection v1.1. Bootstrap values are represented by the size of green circles. Clades with an average branch length distance below one are collapsed. Figure created with iTOL.
