## Additional file 13 for "Automatic annotation of the bHLH gene family in plants"

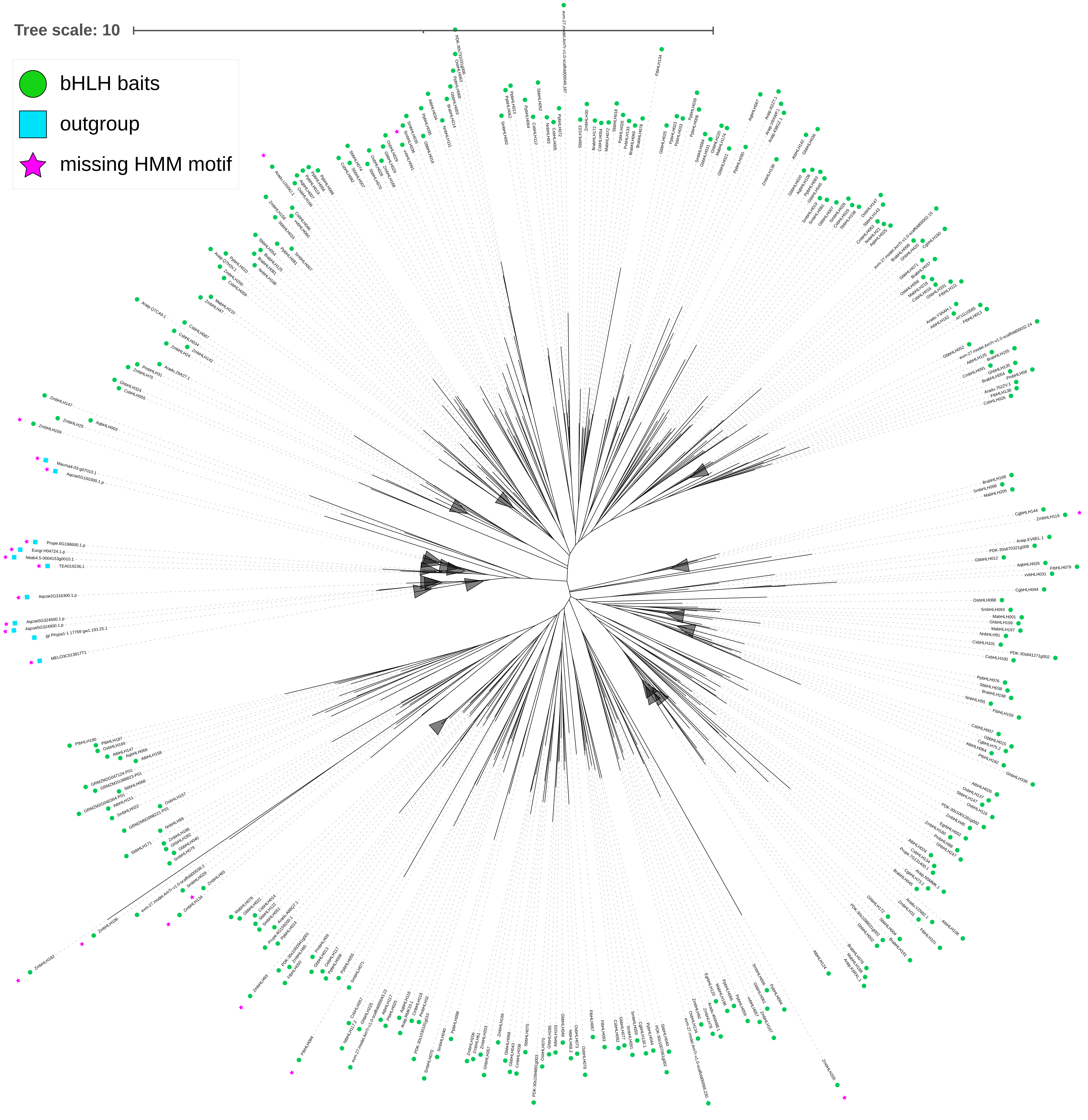


Figure S13: Maximum likelihood tree created with IQ-TREE 1.6.12 showing the phylogenetic relationship between optimised bHLH bait collection and optimised outgroup collection v1.1. Clades with an average branch length distance below one are collapsed. Figure created with iTOL.
